# Characterization of carbon metabolism in a highly adhesive bacterium *Acinetobacter* sp. Tol 5 capable of assimilating diverse hydrocarbons and aromatic compounds

**DOI:** 10.1101/2025.10.30.684093

**Authors:** Shori Inoue, Shogo Yoshimoto, Maiko Hattori, Shotaro Yamagishi, Katsutoshi Hori

## Abstract

Sustainable bioproduction requires developing robust microbial chassis with broad metabolic versatility and suitability for industrial applications. *Acinetobacter* sp. Tol 5 is a highly adhesive bacterium capable of utilizing various hydrocarbons, making it a promising chassis candidate for immobilized whole-cell catalysis. In this study, we characterized the carbon metabolism of Tol 5 by reconstructing metabolic pathway maps from its genomic data and analyzing the transcriptomes of cells grown on ethanol, hexadecane, toluene, and phenol. Genomic analysis revealed that Tol 5 has limited capacity for sugar utilization but possesses a wide range of metabolic pathways for alkane and aromatic compounds, including five distinct aromatic degradation routes that expand the known metabolic diversity of the genus *Acinetobacter*. Transcriptome analysis identified the specific pathway genes induced in response to each carbon source and revealed substrate-dependent cross-regulation between aromatic degradation pathways. Gene disruption experiments further demonstrated that toluene dioxygenase facilitates rapid entry into exponential growth on phenol but reduces carbon assimilation efficiency, while phenol monooxygenase serves as the primary and indispensable route for phenol assimilation, revealing a different physiological role for toluene dioxygenase in Tol 5 compared with *Pseudomonas* putida strains. These findings provide a comprehensive view of the carbon metabolism of Tol 5 and highlight its potential as a microbial chassis for bioprocesses utilizing non-sugar feedstocks, while revealing new aspects of metabolic versatility in the genus *Acinetobacter*.

## 1 Introduction

In recent years, there has been a growing interest in developing sustainable bioprocesses that can utilize diverse carbon sources, including renewable feedstocks and underutilized industrial by-products, to reduce reliance on fossil-derived feedstocks. Microorganisms, especially bacteria, are considered useful hosts for bioconversion due to their rapid growth, ease of genetic manipulation, and capacity for metabolic engineering to improve productivity. *Escherichia coli* is one of the most widely used microbial chassis in bioconversion, owing to its well-characterized molecular biology and metabolic systems (Pontrelli et al., 2018). However, the range of substrates that *E. coli* can natively convert is largely limited to sugars and organic acids, and it has low tolerance to toxic substrates such as aromatic hydrocarbons, which restricts its application in bioprocesses utilizing such compounds. To address this limitation, environmental microorganisms capable of assimilating non-sugar carbon substrates are increasingly valued for converting underutilized resources, including petrochemical by-products, agricultural waste, lignocellulosic biomass, and exhaust gases. For instance, *Pseudomonas putida* is recognized for its high organic solvent tolerance and versatile catabolic pathways, making it particularly suited for bioprocesses involving toxic aromatic substrates (Nakazawa, 2002; Nelson et al., 2002; Ramos et al., 2015; de Lorenzo et al., 2024; Martínez-García and de Lorenzo, 2024). It has been engineered for the production of value-added chemicals from aromatic compounds, including *cis*,*cis*-muconate from lignin-derived substrates (Liu et al., 2024), vanillin from ferulic acid (Ruhl et al., 2025), and medium-chain-length polyhydroxyalkanoates (mcl-PHA) from *p*-coumaric acid (Chen et al., 2025). Collectively, these examples highlight the potential of environmental isolates as microbial chassis for sustainable bioproduction from non-sugar feedstocks and emphasize the importance of exploring and developing new chassis strains capable of metabolizing diverse substrates, tolerating harsh environmental conditions, and performing robustly in industrial bioprocesses.

Members of the genus *Acinetobacter* are ubiquitous, occurring in diverse environments such as soil, marine and freshwater habitats, sediments, and activated sludge (Biggs et al., 2020; Luo et al., 2022). Their extensive genomic plasticity and high tolerance to environmental stress have attracted considerable interest in environmental microbiology. Indeed, several *Acinetobacter* species have shown potential for applications in bioproduction and bioremediation. For example, *Acinetobacter baylyi* ADP1 has been studied as a host for the valorization of aromatic compounds via the β-ketoadipate pathway (Biggs et al., 2020; Luo et al., 2022) and has been engineered for the bioproduction of value-added compounds such as naringenin from *p*-coumaric acid (Kurnia et al., 2024) and mevalonate from lignin-derived substrates (Arvay et al., 2021). *Acinetobacter venetianus* RAG-1, which has multiple alkane hydroxylase genes, has a strong capacity for crude oil degradation and has been explored for bioremediation applications (Liu et al., 2021). Furthermore, *Acinetobacter junii* BP25 has been utilized for the production of polyhydroxybutyrate (PHB) from volatile fatty acids derived from food waste, while *A. venetianus* AMO1502 has been applied to the production of bioemulsifiers (Anburajan et al., 2019; D’Almeida et al., 2024).

### Acinetobacter

sp. Tol 5, originally isolated as a toluene-degrading bacterium from an exhaust gas treatment reactor, can utilize a broad range of non-sugar carbon sources, including aromatic hydrocarbons such as benzene, toluene, and xylene (BTX), as well as carboxylic acids, alcohols, monoterpenes, and triacylglycerols (Hori et al., 2001; Hori et al., 2011; Usami et al., 2018). In addition to this metabolic versatility, Tol 5 exhibits rapid autoagglutination and high adhesiveness to various solid materials, such as plastics, glass, and metals (Ishikawa et al., 2012b). This adhesive phenotype is mediated by its fibrous cell surface protein AtaA, a member of the trimeric autotransporter adhesin (TAA) family (Ishikawa et al., 2012a). Previous studies have revealed the functional domains and biophysical properties of AtaA (Yoshimoto et al., 2023; Yoshimoto et al., 2024) and established a reversible cell immobilization method that enables efficient and resilient attachment of cells to solid carriers (Yoshimoto et al., 2017). Utilizing this immobilization capability, Tol 5 cells expressing heterologous biosynthetic genes have been applied to bioconversion processes, including the gas-phase production of the high-value monoterpenoid (*E*)-geranic acid (Usami et al., 2020). Furthermore, the recent development of CRISPR-Cas9-based genome editing tools for Tol 5 has improved its genetic tractability (Ishikawa and Hori, 2024). Together, the combination of broad substrate range, tolerance to toxic hydrocarbons, AtaA-mediated immobilization capability, and increasing genetic tractability positions Tol 5 as a uniquely promising microbial chassis for bioproduction utilizing non-sugar feedstocks. However, its metabolic network remains largely uncharacterized, limiting the rational engineering of this strain for industrial applications.

A previous study identified a toluene degradation pathway in Tol 5, the first such pathway to be reported in the genus *Acinetobacter*, demonstrating that Tol 5 possesses a broader repertoire for aromatic compound metabolism than is typically found in this genus (Yoshimoto et al., 2025). In this study, we aimed to characterize the carbon metabolism of Tol 5 more comprehensively by reconstructing metabolic pathways based on its complete genome sequence and performing transcriptomic analyses of cells grown on ethanol, hexadecane, or selected aromatic compounds as sole carbon sources.

## 2 Materials and methods

### 2.1 Genome sequence data analysis

The complete genome sequence of Tol 5, consisting of a chromosome and a plasmid, was obtained from the NCBI database (accession: AP024708 and AP024709). All amino acid sequences of the coding sequences (CDSs) were searched against the UniProtKB/Swiss-Prot database (as of August 2024) using BLASTP, and the best-scoring hit for each CDS was used for functional annotation. The sequences were further searched against the KEGG database using BlastKOALA and KEGG automatic annotation server (KAAS) (accession: T11084). For CDSs that could not be annotated with the KEGG database, additional searches were performed against the EggNOG database using eggNOG-mapper. Based on the annotations obtained, a central carbon metabolic pathway map was reconstructed using KEGG mapper. Peripheral metabolic pathways, including those involved in the degradation of alkanes and aromatic compounds, were inferred by comparing operon organization with those previously characterized in closely related bacterial species. Operon structures were predicted using two tools: Operon-mapper (Taboada et al., 2018), which infers operons based on intergenic distances and gene functions, and Rockhopper (McClure et al., 2013), which additionally incorporates transcriptomic data. Functional annotations, operon predictions, and metabolic pathways were manually curated based on information from relevant literature.

### 2.2 Strains, plasmids, and culture conditions

The primers, bacterial strains, and plasmids used in this study are shown in Tables S1 and S2. *E. coli* and its transformants were cultured in lysogeny broth (LB) medium (20066-95; Nacalai Tesque, Kyoto, Japan) at 37°C. *Acinetobacter* sp. Tol 5 and its mutants were cultured in LB medium or basal salt (BS) medium (Hori et al., 2001) at 28°C.

### 2.3 Gene knockout

The gene knockout mutants were generated using the cytidine base editing system for *Acinetobacter* according to the previous reports (Wang et al., 2019; Inoue et al., 2025). The DNA dimer fragment encoding the sequence of the single guide RNA was prepared by mixing oligo DNAs listed in Table S2 in 50 mM NaCl solution and gradually cooling from 95°C to 18°C at 0.1°C/s. This dimer was introduced into the BsaI site of pBECAb-apr. The constructed plasmid was electroporated into Tol 5REK, a restriction-modification system and *ataA*-deficient strain (Ishikawa and Hori, 2024). After overnight culture of Tol 5REK harboring the plasmid in BS medium to promote mutation, the cells were spread on a BS agar plate containing 5 % (w/v) sucrose to remove the plasmid. Mutation of the target gene was confirmed by sequencing the DNA amplified from the genome by PCR using KOD FX Neo (Toyobo, Osaka, Japan).

### 2.4 Growth assay on various carbon sources

The Tol 5 Δ*ataA* mutant cells cultured in 5 mL of LB medium at 28°C for 24 hours were collected by centrifugation at 8000 ×*g* for 5 min at room temperature, washed with 5 mL of BS medium, and resuspended in BS medium at OD660 = 2.0. For agar culture, the cell suspension was streaked onto BS agar plates containing each carbon source at a carbon equivalent concentration of 1.4 × 10^-2^ mol/L. Each plate was individually sealed in an AnaeroPack pouch (A-65; Mitsubishi Gas Chemical, Tokyo, Japan) to prevent contamination with volatile compounds from the surrounding environment and incubated at 28°C for 5 days. For volatile substrates, 10 µL of the compound was absorbed onto a tissue paper and placed inside the pouch together with the plate before sealing.

For liquid culture, a 250-µL aliquot of the suspension was inoculated into 5 mL of BS medium in a glass test tube. Each carbon source was added at a carbon equivalent concentration of 1.4 × 10^-2^ mol/L. The test tubes were then capped with silicone stoppers for non-volatile substrates or viton rubber stoppers for volatile substrates, and incubated at 28°C with shaking at 200 rpm. The optical density (OD) was monitored every 60 min using the OD-monitorC&T system (Taitec, Saitama, Japan).

### 2.5 Growth assay on *n*-alkanes of various chain lengths

The Tol 5 Δ*ataA* mutant cells cultured in 5 mL of LB medium at 28°C for 24 hours were collected by centrifugation at 8000 ×*g* for 5 min at room temperature, washed with 5 mL of BS medium, and resuspended in BS medium at OD660 = 2.0. For solid long-chain alkanes (tetracosane and dotriacontane), the compounds were first dissolved in chloroform, added to 100-mL Erlenmeyer flasks, and the solvent was allowed to evaporate completely before use. To each flask, 20 mL of BS medium, four rectangular pieces of polyurethane foam support (10 × 10 × 10 mm, CFH-20, 20 pores/25 mm; Inoac Corporation, Nagoya, Japan) to aid mixing of solid or oil alkanes, and 200 µL of the cell suspension were added. Liquid alkanes (dodecane and hexadecane) were added to each flask at a carbon equivalent concentration of 1.4 × 10⁻² mol/L. Flasks were capped with silicone stoppers and incubated with shaking at 115 rpm and 28°C. At each time point, a 500-µL aliquot was withdrawn from each flask and the OD at 660 nm (OD660) was measured using a UV‒Vis spectrophotometer (UV-1850; Shimadzu Corporation, Kyoto, Japan). For octane and dodecane, a 50-µL aliquot of the suspension was inoculated into 5 mL of BS medium in a glass test tube. Each alkane was added at a carbon equivalent concentration of 1.4 × 10^-2^ mol/L. The test tubes were then capped with viton rubber stoppers for volatile substrates, and incubated at 28°C with shaking at 200 rpm. The OD was monitored every 60 min using the OD-monitorC&T system.

### 2.6 Growth assay of Δ*todC1*, Δ*todE*, and Δ*mphN* mutants

The Tol 5REK and its derivative mutant cells were cultured in 5 mL of LB medium supplemented with each carbon source at a carbon equivalent concentration of 1.4 × 10^-2^ mol/L at 28°C for 24 hours to induce the relevant catabolic genes. The cells were then collected by centrifugation at 8000 ×*g* for 5 min at room temperature, washed with 5 mL of BS medium, and resuspended in BS medium at OD660 = 2.0. A 250-µL aliquot of the suspension was inoculated into 5 mL of BS medium in a glass test tube. Each carbon source was added at a carbon equivalent concentration of 1.4 × 10^-2^ mol/L. The test tubes were capped with silicone stoppers for non-volatile substrates or Viton rubber stoppers for volatile substrates. The OD was monitored every 60 min using the OD-monitorC&T system.

### 2.7 Bacterial culture conditions for transcriptome analysis

The wild type strain of Tol 5 was cultured in 5 mL of LB medium supplemented with each carbon source at a carbon equivalent concentration of 3.3 × 10^-2^ mol/L at 28°C for 24 hours. The cells were then collected by centrifugation at 8000 ×*g* for 5 min at room temperature, washed with 5 mL of BS medium, and resuspended in BS medium at OD660 = 2.0. A 200-µL aliquot of the suspension was inoculated into 20 mL of BS medium in 100-mL Erlenmeyer flasks. Because Tol 5 exhibits strong autoagglutination and high adhesiveness mediated by the cell surface protein AtaA, cells cultured in BS medium do not remain in suspension but instead form aggregates and adhere to surfaces; therefore, four rectangular pieces of polyurethane foam support (CFH-20) were added to each flask as carriers to immobilize the cells. Each carbon source was added at a carbon equivalent concentration of 3.3 × 10^-2^ mol/L. The flasks were capped with silicone stoppers for cultures containing sodium lactate or hexadecane, or with Viton rubber stoppers for those containing ethanol, toluene, or phenol. The cultures were incubated at 28°C with shaking at 115 rpm. At each time point, flasks were collected, and the cultures were mixed vigorously with 1 mL of 10% Casamino Acids technical grade (CA-T; Becton, Dickinson and Company, Franklin Lakes, NJ, USA) to inhibit AtaA-mediated cell aggregation and to detach the bacterial cells from the support, as previously reported (Ohara et al., 2019). The OD₆₆₀ of the resulting cell suspension was measured using the UV–Vis spectrophotometer. Cells were harvested from the polyurethane foam support at OD₆₆₀ = 0.2–0.5 for RNA extraction.

### 2.8 RNA sequencing

Cells were collected by centrifugation (8,000 × g, 4°C, 3 min), and the cell pellets were stored at - 80°C. For each growth condition, three biological replicates were prepared. Total RNA was extracted using the RNA Prep Kit (Cica Genus, Tokyo, Japan) according to the manufacturer’s protocol. Ribosomal RNA was removed using the NEBNext rRNA Depletion Kit (Bacteria) (New England Biolabs, Ipswich, MA, USA), and cDNA libraries were generated with the NEBNext Ultra II RNA Library Prep Kit for Illumina (New England Biolabs, Ipswich, MA, USA). The cDNA libraries were sequenced on the Illumina NextSeq550. Raw FASTQ reads were quality-filtered using fastp (version 0.23.2) and mapped to the Tol 5 genome (accession: AP024708 and AP024709) using Bowtie2 (version 2.5.1) with default parameters. Mapped reads were counted using featureCounts (version 2.0.6).

### 2.9 Differential gene-expression analysis

Raw read counts of CDSs were analyzed in R using the edgeR package (version 3.42.4). Genes with extremely low expression were removed using the filterByExpr function. Normalization was performed using the trimmed mean of M-values (TMM) method implemented in the calcNormFactors function. Differential gene expression analysis was conducted using the quasi-likelihood pipeline. A design matrix was constructed incorporating the cells grown on six different carbon sources (n = 3), with the data from those grown on lactate set as the reference group. Dispersion estimates were obtained using the estimateDisp function, and model fitting was performed with the glmQLFit function. Differential expression between each condition and the control condition (lactate condition) was assessed using the glmQLFTest function. Genes with false discovery rate (FDR) < 0.01, |log2 fold change (FC)| > 1, and log2 counts per million (CPM) > 3 were extracted as significantly differentially expressed genes (DEGs). Over-representation analysis (ORA) was performed separately on upregulated and downregulated DEGs under each condition using the enricher function in the clusterProfiler package (version 4.12.6) in R (Yu et al., 2012), with KEGG pathway annotations as the gene set database and all KO identifiers detected in the Tol 5 genome as the background gene set. Enrichment was assessed using the Benjamini–Hochberg method for *p*-value adjustment, with minimum and maximum gene set sizes set to 5 and 5,000, respectively. The top 10 pathways ranked by gene count were selected for each direction and visualized as bar plots. All figures were generated using ggplot2 (version 3.5.2) in R.

## 3 Results

### 3.1 Genomic features of central carbon metabolism in Tol 5

The complete genome of Tol 5 consists of two circular DNA components, a chromosome of 4,681,789 bp and a plasmid of 117,717 bp, collectively encoding 4,404 predicted CDSs (Ishikawa and Hori, 2021). To predict the metabolic potential of Tol 5, we first evaluated the functional annotation of each CDS using multiple databases, including NCBI, UniProtKB/Swiss-Prot, KEGG, and EggNOG, as well as information from relevant literature (Table S3). Tol 5 encodes all the enzymes for gluconeogenesis, pentose phosphate pathway, tricarboxylic acid (TCA) cycle, glyoxylate shunt, purine and pyrimidine biosynthesis, β-oxidation, and fatty acid biosynthesis (Fig. 1, Table S4). In contrast, glycolysis via the Embden–Meyerhof–Parnas (EMP) pathway is incomplete due to the absence of hexokinase, phosphofructokinase, and pyruvate kinase. It also lacks the genes for the Entner–Doudoroff (ED) pathway, an alternative glucose utilization route, and a pentose utilization pathway encoded by *ara* genes that converts pentoses into α-ketoglutarate. To confirm this limited sugar utilization capacity, we tested the growth of Tol 5 on agar plates supplemented with D-glucose, D-fructose, L-arabinose, or sodium gluconate as sole carbon sources and observed no growth under any of these conditions (Fig. S1). These results suggest that Tol 5 has a very limited capacity for sugar utilization as a carbon source.

**Fig. 1.**
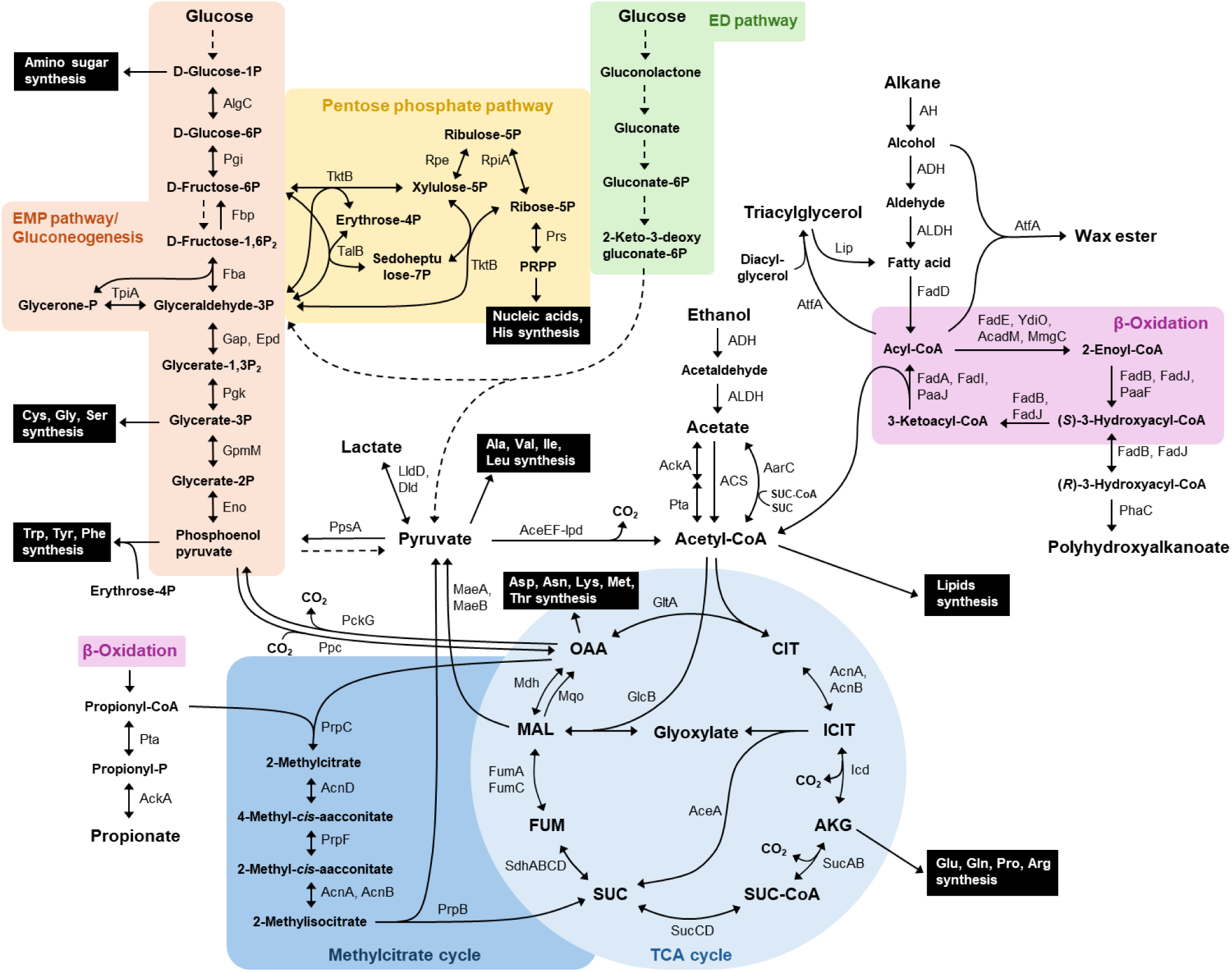
Prediction of the central metabolic pathway of *Acinetobacter* sp. Tol 5. Genes related to the central metabolic pathway were analyzed and listed in Table S4. Dashed arrows indicate metabolic reactions for which the corresponding genes are not found in Tol 5 genome. Abbreviations: ADH; alcohol dehydrogenase, AH; alkane hydroxylase, AKG; α-ketoglutarate, ALDH; aldehyde dehydrogenase, CIT; citrate, FUM; fumarate, ICIT; isocitrate, MAL; malate, OAA; oxaloacetate, PRPP; phosphoribosyl pyrophosphate, SUC; succinate, SUC-CoA; succinyl-CoA.

As with many *Acinetobacter* strains (Ren and Palmer Lauren, 2023), alcohols, carboxylic acids, and fatty acids serve as carbon sources for Tol 5. The genome encodes multiple putative alcohol dehydrogenase (ADH) and aldehyde dehydrogenase genes (Table S4), suggesting a capacity to oxidize alcohols to their corresponding carboxylic acids. Three acetate assimilation routes were identified, including the reversible acetate kinase-phosphotransacetylase (AckA-Pta) pathway, the irreversible acetyl-CoA synthetase (ACS) pathway, and the reversible pathway mediated by succinyl-CoA:coenzyme A transferase AarC (Fig. 1, Table S4). For fatty acid degradation, multiple candidate genes corresponding to each step of β-oxidation were identified, including two putative acyl-CoA ligases, twelve acyl-CoA dehydrogenases, four enoyl-CoA hydratases, and three thiolases. Propanoyl-CoA, generated during β-oxidation of odd-chain fatty acids, was predicted to be catabolized to succinate and pyruvate via the methylcitrate cycle, which consists of 2-methylisocitrate lyase (PrpB), 2-methylcitrate synthase (PrpC), 2-methylcitrate dehydratase (AcnD), 2-methylaconitate isomerase (PrpF), and bifunctional aconitate hydratase/isomerase (AcnA and/or AcnB). In addition, four extracellular lipase genes were identified in Tol 5 (Table S4), all of which are tandemly arranged (*TOL5_03740*–*TOL5_03770*) and share 44–52% amino acid sequence identities with the triacylglycerol lipase Lip reported in *P. aeruginosa* PAO1 (Wohlfarth et al., 1992). Tol 5 further encodes key enzymes that redirect fatty acid oxidation intermediates into carbon storage compounds. A bifunctional wax ester synthase/acyl-CoA:diacylglycerol acyltransferase (AtfA) mediates the production of wax esters and triacylglycerols, while polyhydroxyalkanoate (PHA) synthase (PhaC) is responsible for the production of PHAs.

### 3.2 Genomic features of alkane metabolism in Tol 5

Under aerobic conditions, *n*-alkanes are first oxidized at their terminal methyl group to primary alcohols by alkane hydroxylases, and these alcohols are then further oxidized to aldehydes and finally to fatty acids (Rojo, 2009). The Tol 5 chromosome encodes multiple alkane hydroxylases, including a membrane-bound non-heme iron alkane monooxygenase AlkB, and three flavin-containing monooxygenases encoded by *almA*, *ladA_1*, *ladA_2* (Table S4). Additionally, the plasmid of Tol 5 carries a gene encoding an alkane hydroxylase (*TOL5_43480*), whose amino acid sequence shares 99% identity with that of a cytochrome P450 monooxygenase of the CYP153 family found in *Acinetobacter* sp. EB104 (Maier et al., 2001) along with its associated redox partners, ferredoxin (*TOL5_43490*) and ferredoxin reductase (*TOL5_43470*). AlkB and CYP153 typically oxidize short-and medium-chain *n*-alkanes (C16 or less), whereas AlmA and LadA act on long-chain *n*-alkanes (C20 or more) (Nie et al., 2014; Kong et al., 2021; Chen et al., 2024). To evaluate the alkane utilization capacity of Tol 5, we examined growth on *n*-alkanes ranging from C8 to C32 as sole carbon sources (Fig. S2). Tol 5 grew on dodecane (C12), hexadecane (C16), tetracosane (C24), and dotriacontane (C32), but not on octane (C8), demonstrating that Tol 5 can utilize *n*-alkanes of at least C12 and longer as carbon sources. The resulting primary alcohols are further oxidized by alcohol dehydrogenases (ADHs) and aldehyde dehydrogenases (ALDHs). In Tol 5 genome, at least six putative ADH genes and one ALDH gene were predicted to be involved in fatty alcohol and fatty aldehyde oxidation (Table S4). Although the functions of most ADHs in Tol 5 remain unclear, the iron-containing ADH encoded by *yiaY_1* shares 91% amino acid sequence identity with ADH4 from *Acinetobacter baumannii* ATCC 19606, which catalyzes the oxidation of ethanol, 1-propanol, and 1- butanol (Lin et al., 2021). The putative ALDH encoded by *aldB* (AldB) shares 87% amino acid identity with a long-chain ALDH, Ald1, reported in *Acinetobacter* sp. M-1 (Ishige et al., 2000).

### 3.3 Genomic features of aromatic compound metabolism in Tol 5

Genes encoding multi-component oxygenases for aromatic compound oxidation, as well as enzymes for successive reaction steps leading to central metabolites, are generally organized into operons and regulated by specific transcriptional regulators (Phale et al., 2020). Based on the predicted operons involved in aromatic compound degradation (Table S4, Fig. 2), we reconstructed the aromatic degradation pathways in Tol 5, identifying five major aromatic degradation routes (Fig. 3).

**Fig. 2.**
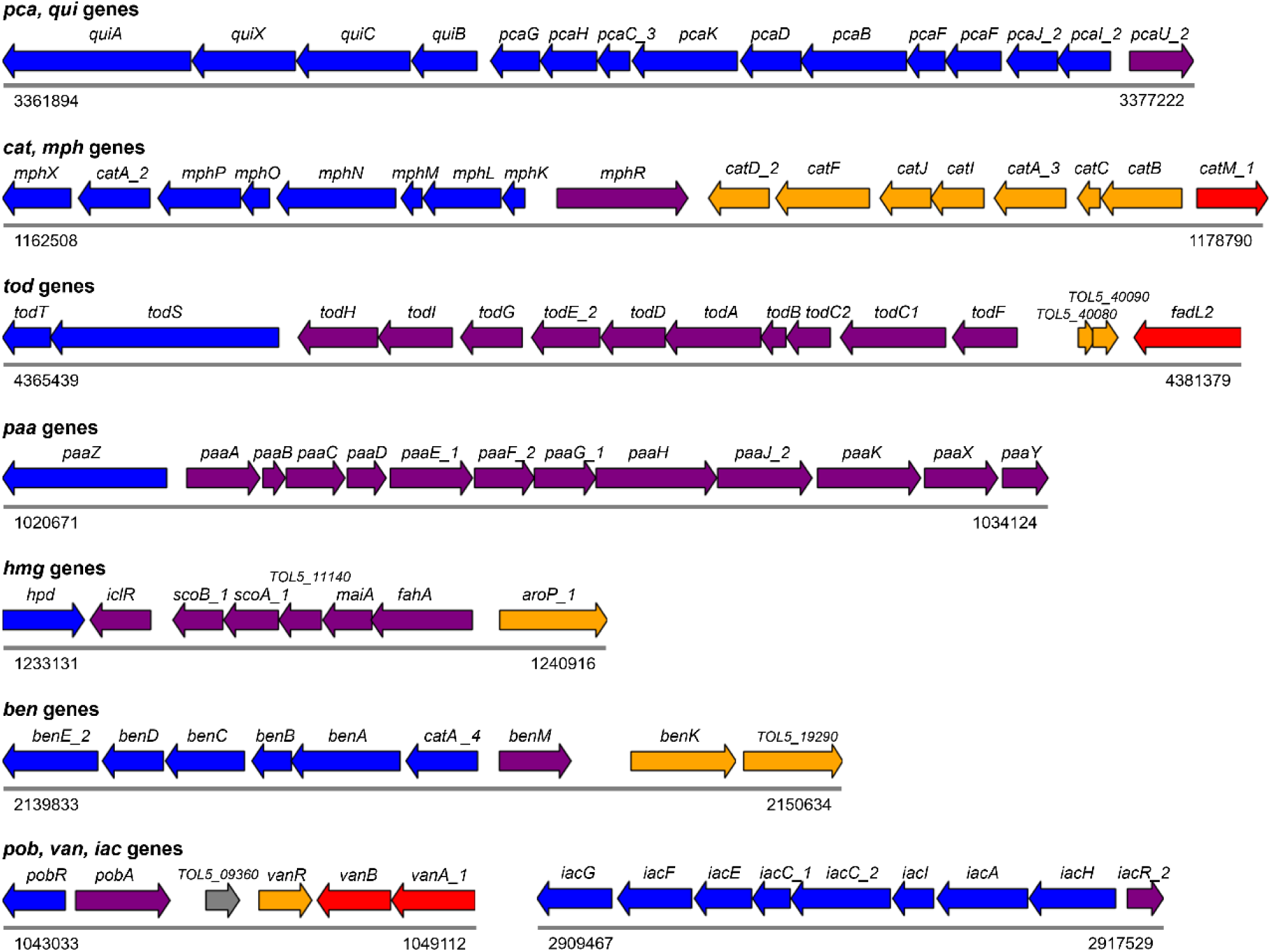
Genetic organization of genes involved in the degradation of aromatic compounds in *Acinetobacter* sp. Tol 5. Predicted operon structures are illustrated based on analyses using Operon-mapper and Rockhopper. Genes within the same operon are represented by arrows of the same color. Proposed gene names or predicted functions are labeled above each arrow, and genomic coordinates are indicated below.

**Fig. 3.**
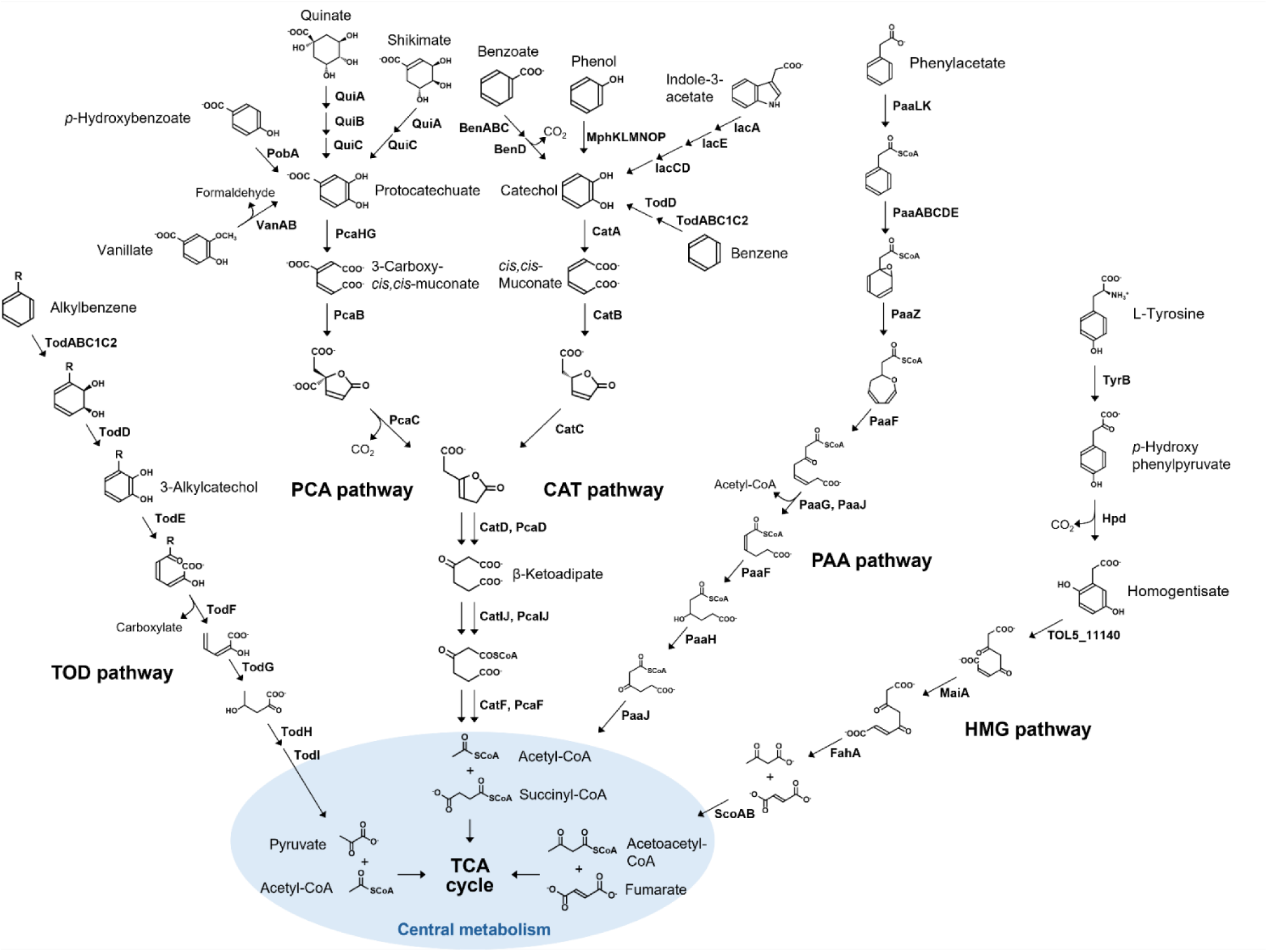
Predicted degradation pathways of aromatic compounds in *Acinetobacter* sp. Tol 5, reconstructed based on the genes shown in Fig. 2 and Table S4. Question marks and dashed arrows indicate steps for which the genes encoding the corresponding enzymes have not been identified in Tol 5.

The *pca* genes encode the enzymes responsible for the degradation of protocatechuate (PCA pathway) (Fig. 3), forming a gene cluster together with the aromatic compound transport protein PcaK and the transcriptional repressor PcaU (Fig. 2). The *cat* gene cluster encodes the enzymes responsible for the degradation of catechol (CAT pathway) (Fig. 3) and the transcriptional activator CatM (Fig. 2). Protocatechuate and catechol are common intermediates derived from various aromatic compounds through various upstream pathways. The PCA and CAT pathways funnel these intermediates into acetyl-CoA and succinyl-CoA via oxidative *ortho*-cleavage of the aromatic ring, serving as central routes for aromatic compound degradation that are widely conserved in *Acinetobacter* species (Barbe et al., 2004; Fischer et al., 2008; Stuani et al., 2014; Breisch et al., 2022). In Tol 5, the *pca* genes are located adjacent to the *qui* genes (Fig. 2), which encode enzymes that convert quinate and shikimate to protocatechuate (Fig. 3). The structure of this gene cluster is similar to that reported in *A. baylyi* ADP1, in which the *pca* and *qui* genes are organized as a large single operon under the regulator PcaU (Dal et al., 2005) (Fig. S3). Tol 5 also possesses the *pob* and *van* genes, which encode upstream pathways leading to protocatechuate, along with their respective transcriptional regulators (PobR and VanR) (Fig. 2 and 3). Consistent with these genomic predictions, Tol 5 grew on quinate, shikimate, *p*-hydroxybenzoate, and vanillate as sole carbon sources in both agar and liquid culture (Fig. S4A and S5A).

The *cat* genes are located adjacent to the *mph* gene cluster, which encodes the components of the phenol monooxygenase complex MphKLMNOP, a catechol 1,2-dioxygenase, and the regulatory proteins MphX (repressor) and MphR (activator) (Fig. 2). Notably, Tol 5 encodes three catechol 1,2-dioxygenase paralogs distributed across distinct operons: CatAmph (encoded by *catA_2*, within the *mph* operon), CatAcat (encoded by *catA_3*, within the *cat* operon), and CatAben (encoded by *catA_4*, within the *ben* operon). The structure of the *mph* operon in Tol 5 resembles that of *Acinetobacter calcoaceticus* PHEA-2 (Yu et al., 2011), but is distinguished by the presence of the additional *catA_2* gene, encoding CatAmph, located immediately downstream within the same operon (Fig. S3). In addition, Tol 5 possesses the *ben* and *iac* genes, which encode enzymes for the upstream pathways leading to catechol, along with their respective transcriptional regulators (BenM and IacR) and transporters (BenE, BenK, and a putative transporter encoded by *TOL5_19290*) (Fig. 2 and 3). The *ben* operon includes *catA_4* encoding CatAben, distinguishing it from those reported in *A. baylyi* ADP1 (Bleichrodt et al., 2010) (Fig. S3). Tol 5 grew on phenol and benzoate as sole carbon sources, supporting the predicted roles of the *mph* and *ben* pathways in the assimilation of these substrates (Fig. S4B and S5A).

The *tod* operon encodes the enzymes responsible for the degradation of alkylbenzenes (TOD pathway) (Fig. 3) and is located between the *todS*-*todT* two-component regulatory system and the putative toluene transporter gene *fadL2*, which is in a separate operon (Fig. 2). In the TOD pathway, alkylbenzenes are first oxidized to 3-alkylcatechols. The aromatic ring is then opened via oxidative *meta*-cleavage, and the intermediates are ultimately converted into carboxylate, pyruvate, and acetyl-CoA. While the TOD pathway is rare within the genus *Acinetobacter*, which has been identified only in *Acinetobacter johnsonii*, *Acinetobacter pittii*, and *Acinetobacter* sp. ANC 4635, the *tod* genes of Tol 5 show high similarity with those of *P. putida* F1 (Yoshimoto et al., 2025). We have previously reported that the toluene dioxygenase complex encoded by todABC1C2 plays a central role in toluene and benzene assimilation in Tol 5 (Yoshimoto et al., 2025). In addition to these substrates, we confirmed that Tol 5 also grew on ethylbenzene as sole carbon sources (Fig. S4C and S5B), suggesting that the TOD pathway supports the assimilation of a broader range of alkylbenzenes.

In addition to pathways for exogenous aromatic compounds, Tol 5 encodes two further aromatic degradation routes likely associated with amino acid catabolism. Tol 5 possesses the *paa* genes, which encode the enzymes responsible for the degradation of phenylacetic acid (PAA pathway) (Fig. 3), as well as the transcriptional repressor PaaX (Fig. 2). The PAA pathway degrades phenylacetate via epoxidation of CoA thioesters. While this pathway in *A. baumannii* strains is thought to be primarily associated with phenylalanine degradation (Teufel et al., 2010; Hooppaw Anna et al., 2022), Tol 5 lacks the enzymes required to convert phenylalanine to phenylacetate. Tol 5 also possesses a gene cluster encoding a putative homogentisate degradation route (HMG pathway) (Fig. 3), which includes 4-hydroxyphenylpyruvate dioxygenase (HPD), maleylacetoacetate isomerase (MaiA), fumarylacetoacetase (FahA), 3-oxoacid CoA-transferase subunits A and B (ScoA and ScoB), a putative homogentisate dioxygenase (*TOL5_11140*), an aromatic amino acid transporter (AroP), and an IclR-type transcriptional repressor (IclR) (Fig. 2). Although TOL5_11140 was annotated as a glyoxalase (Table S3), a recent study demonstrated that this homologous gene in *A. baumannii* functions as a homogentisate 1,2-dioxygenase despite its sequence similarity to glyoxalases (Seo et al., 2025). To evaluate whether Tol 5 can assimilate aromatic amino acid-related compounds as sole carbon sources, we tested its growth on L-phenylalanine and, instead of poorly soluble L-tyrosine, on its downstream intermediates homogentisate and *p*-hydroxyphenylpyruvate (Fig. S4D). However, Tol 5 did not grow on any of these compounds as the sole carbon source. Phenylacetate was not tested as a growth substrate because phenylacetic acid and its salts are subject to regulations as controlled drug precursors, making procurement difficult. Therefore, the physiological roles of the PAA and HMG pathways remain unclear, although these pathways may be involved in intracellular amino acid catabolism rather than serving as routes for exogenous carbon assimilation.

### 3.4 Transcriptomic responses of cells grown on alcohol and alkane carbon sources

While genomic analysis revealed that Tol 5 possesses multiple genes involved in the metabolism of alcohols and alkanes, the expression of key enzymes, such as alcohol dehydrogenases and alkane hydroxylases, is generally subject to strict regulation depending on the available carbon source. To identify the specific genes induced for the assimilation of each substrate, we performed transcriptome analysis using lactate, ethanol, and hexadecane as sole carbon sources. Lactate was selected as the reference condition because it supports robust exponential growth of Tol 5 (Hori et al., 2011) and enters central carbon metabolism via pyruvate and the TCA cycle, without requiring the peripheral oxidation steps involved in alcohol or alkane assimilation. Ethanol and hexadecane were chosen as representative substrates for alcohols and alkanes, respectively, as both are widely used model compounds for studying alcohol and alkane catabolism. Tol 5 cells were cultured in BS medium supplemented with each carbon source and their transcriptomes were compared with those of cells grown on lactate (Tables S5 and S6). DEGs were defined by the thresholds of |log2 FC| > 1, log2 CPM > 3, and FDR < 0.01.

In the ethanol culture, 14 genes were significantly upregulated, and 57 genes were downregulated (Fig. 4A, Table S7). The limited number of upregulated DEGs in ethanol culture likely reflects the fact that ethanol assimilation requires only a small number of dedicated enzymes for its conversion to acetyl-CoA. Among the 14 upregulated genes, pathways related to central carbon metabolism were significantly enriched, including glycolysis/gluconeogenesis, pyruvate metabolism, and the citrate cycle (Fig. 4B). These categories include the ethanol assimilation genes described below. In contrast, downregulated genes were overrepresented in pathways for the degradation of branched-chain amino acids, fatty acids, and benzoate, suggesting that growth on ethanol as the sole carbon source represses the expression of genes involved in alternative substrate utilization. Focusing on the ethanol assimilation pathway, among the upregulated genes, *yiaY_1* and *aldB*, which encode an iron-containing ADH and a putative ALDH, respectively, were identified (Table S7). Since *yiaY_1* was the only ADH gene induced by ethanol, it is likely the primary ADH responsible for ethanol oxidation in Tol 5 (Fig. 4C). Although AldB shares 87% amino acid identity with Ald1 from *Acinetobacter* sp. M-1, which has been characterized as a long-chain ALDH that is inactive toward short-chain substrates such as acetaldehyde (Ishige et al., 2000), its induction during growth on ethanol suggests that AldB functions differently from Ald1 and catalyzes acetaldehyde oxidation in Tol 5. Regarding the conversion of acetate into acetyl-CoA, although Tol 5 possesses both the AckA-Pta pathway and the ACS pathway (Fig. 1), genes involved in neither pathway were significantly upregulated (Fig. 4C).

**Fig. 4.**
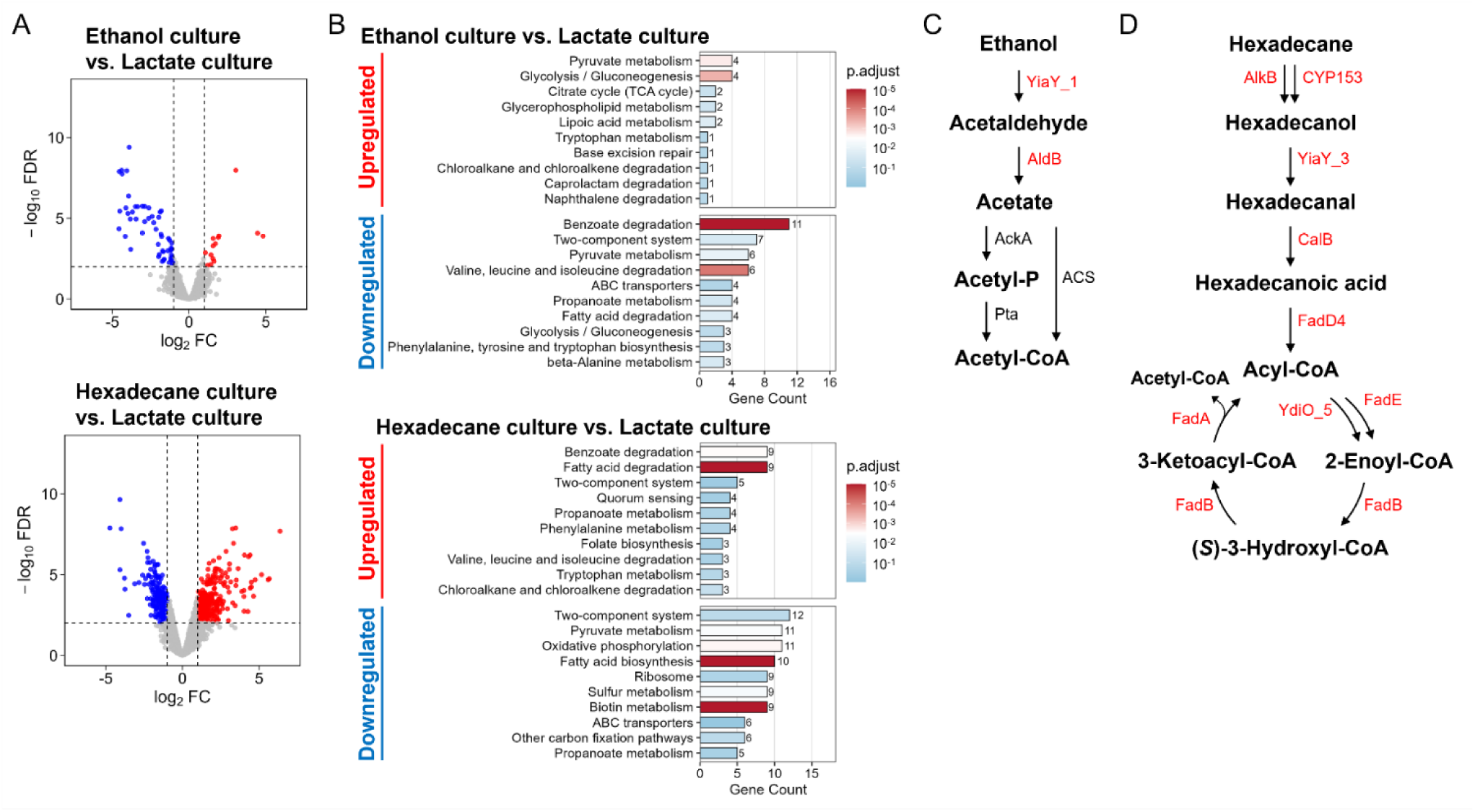
Transcriptome analysis of *Acinetobacter* sp. Tol 5 grown on ethanol, and hexadecane using lactate as the reference condition. (**A**) Volcano plots comparing gene expression between lactate and ethanol or hexadecane as sole carbon sources (n = 3 biologically independent samples). Red and blue dots represent significantly upregulated and downregulated genes, respectively (FDR < 0.01, |log₂ FC| > 1). (**B)** Bar plots showing the results of ORA based on KEGG pathway for upregulated and downregulated DEGs in ethanol and hexadecane cultures. The top 10 pathways ranked by gene count are shown for each direction. Bar length represents the number of DEGs annotated to each pathway, and bar color indicates the adjusted *p*-value (p.adjust). **(C**, **D**) Proposed ethanol (**C**) and hexadecane (**D**) degradation pathways in Tol 5. Solid arrows indicate the primary metabolic pathways predicted from transcriptome data. Genes encoding enzymes significantly upregulated under each condition are shown in red.

In the hexadecane culture, 266 genes were significantly upregulated and 196 genes were downregulated (Fig. 4A, Table S7). Among the upregulated genes, fatty acid degradation was the most significantly enriched pathway, followed by benzoate degradation (Fig. 4B). In contrast, downregulated genes were predominantly characterized by enrichment of fatty acid biosynthesis and biotin metabolism, suggesting a metabolic shift away from lipid biosynthesis under conditions where hexadecane serves as the sole carbon source. Focusing on the hexadecane assimilation pathway, among the multiple alkane hydroxylase genes in Tol 5, *alkB* and the CYP153 gene were specifically induced, indicating their involvement in the initial hydroxylation of hexadecane (Fig. 4D). The genes *yiaY_3*, encoding an iron-containing ADH, and *calB*, encoding a putative coniferyl aldehyde dehydrogenase, were also upregulated, suggesting their roles in the oxidation of hexadecanol to hexadecanal and subsequently to hexadecanoic acid. The genes involved in every step of the β-oxidation were upregulated. Among the multiple predicted enzymes for each step, only *fadA*, *fadB*, *fadD4* (one of the two predicted acyl-CoA ligase genes), *fadE*, and *ydiO_5* (two of the twelve predicted acyl-CoA dehydrogenase genes) were specifically induced (Fig. 4D).

### 3.5 Transcriptomic responses of cells grown on aromatic carbon sources

Subsequently, we focused on the transcriptional regulation of the aromatic degradation pathways in Tol 5, particularly for the degradation of toluene and phenol. Although both compounds are recalcitrant and highly toxic to microbial cells, Tol 5 exhibited relatively rapid growth on these substrates as sole carbon sources (Fig. S5A and B). Tol 5 cells were cultured in BS medium supplemented with toluene or phenol and subjected to transcriptome analysis using the same procedure described above. The resulting data were also compared with those of cells grown on lactate.

In the toluene and phenol cultures, 349 and 643 genes were upregulated, while 292 and 578 genes were downregulated, respectively (Fig. 5A, Table S7). Under both conditions, multiple pathways associated with aromatic compound degradation were significantly enriched among the upregulated genes, including benzoate degradation, toluene degradation, xylene degradation, and chlorocyclohexane and chlorobenzene degradation (Fig. 5B). The enrichment of chloroalkane and chloroalkene degradation pathways likely reflects the induced aromatic oxygenases, some of which are known to act on both aromatic and halogenated hydrocarbons. Biosynthesis of various siderophores was also enriched in the toluene culture, indicating a response to an increased demand for iron acquisition to support the activity of aromatic oxygenases, which require iron as a cofactor. In addition to these responses, fatty acid degradation and phenylalanine metabolism were enriched exclusively in the toluene culture, suggesting that toluene assimilation involves a broader remodeling of peripheral catabolic pathways compared to those in lactate culture. In contrast, the downregulated genes were most prominently enriched in ribosomal components under both conditions. Pyruvate metabolism and oxidative phosphorylation were also enriched, suggesting that central carbon and energy metabolism was altered during growth on the aromatic substrates compared with growth on lactate.

**Fig. 5.**
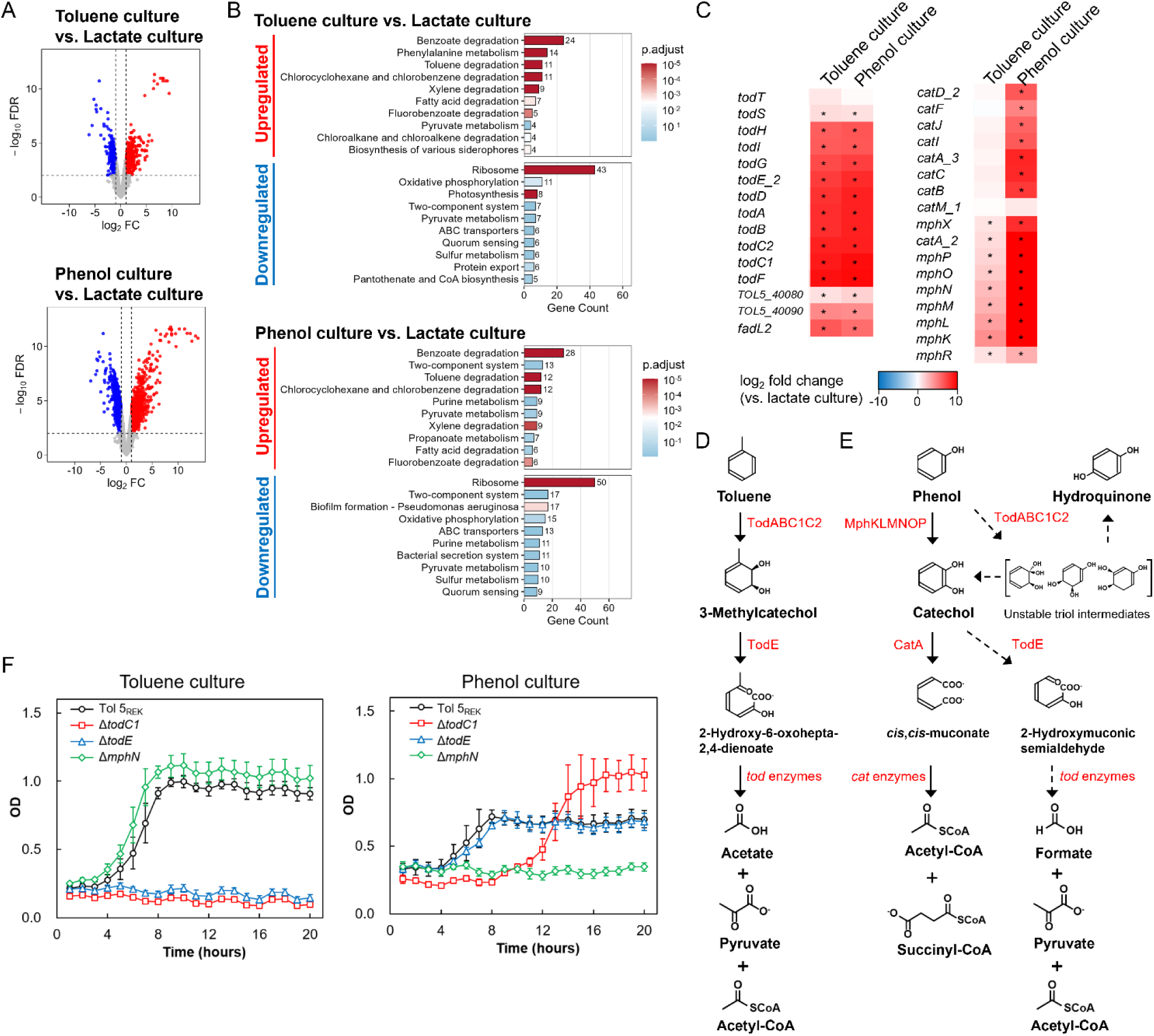
Differential gene expression of *Acinetobacter sp.* Tol 5 grown on toluene and phenol using lactate as the reference condition. (**A**) Volcano plots comparing gene expression between lactate and toluene or phenol as sole carbon sources (n = 3 biologically independent samples). Red and blue dots represent significantly upregulated and downregulated genes, respectively (FDR < 0.01, |log₂ FC| > 1). (**B)** Bar plots showing the results of ORA based on KEGG pathway for upregulated and downregulated DEGs in toluene and phenol cultures. The top 10 pathways ranked by gene count are shown for each direction. Bar length represents the number of DEGs annotated to each pathway, and bar color indicates the adjusted *p*-value (p.adjust). (**C**) Heat maps showing the average log₂ FC of the *tod*, *mph*, and *cat* genes. The color gradient indicates the magnitude of the log₂ FC values, with the deepest red representing ≥ 10 and the deepest blue representing ≤-10. Asterisks (*) denote differentially expressed genes defined by |log2 FC| > 1, log2 CPM > 3, and FDR < 0.01. (**D, E**) Proposed degradation pathways of toluene (**D**) and phenol (**E**) in Tol 5. Solid arrows indicate the primary metabolic pathways predicted from transcriptome data. Dashed arrows indicate pathways that were induced at the transcriptional level but whose contribution to substrate assimilation was not confirmed by gene disruption experiments. Enzyme names shown in red indicate significantly upregulated genes under each condition. (**F**) Growth of gene disruption mutants on toluene and phenol. Tol 5REK (circles), the Δ*todC1* mutant (squares), the Δ*todE* mutant (triangles), and the Δ*mphN* mutant (diamonds) were inoculated to BS medium containing toluene or phenol at a carbon equivalent concentration of 1.4 × 10^-^² mol/L, and OD was monitored using the OD-monitorC&T system. Data are presented as the means ± SEMs (biological replicates n = 3).

Regarding the specific degradation pathways induced for each substrate, the toluene culture strongly induced the *tod* operon, including all of the TOD pathway genes (Fig. 5C). Although genes in the *mph* operon (encoding phenol monooxygenase) and the *paa* operon (encoding the PAA pathway) showed slight upregulation, their induction levels were negligible compared to those of the *tod* operon, and most other aromatic degradation pathway genes were not significantly changed (Fig. 5C and S6). The *fadL2*, which is located adjacent to the *tod* operon and encodes a putative toluene transporter (Fig. 2), was also upregulated (Fig. 5C). Consistent with these transcriptional responses and previous study showing that disruption of *todC1* abolishes growth on toluene (Yoshimoto et al., 2025), these results indicate that toluene degradation proceeds exclusively through the TOD pathway (Fig. 5D). In contrast, in the phenol culture, both the *mph* and *cat* operons, which comprise the predicted phenol degradation pathway, and the *tod* operon were significantly upregulated (Fig. 5C). Notably, the induction levels of the *tod* genes reached those observed in toluene culture, suggesting an involvement of *tod* enzymes in phenol metabolism. Most genes of the PCA, PAA, and HMG pathways were not significantly changed in phenol culture (Fig. S6).

Although phenol is predicted to be metabolized via hydroxylation by phenol monooxygenase and the CAT pathway, encoded by the *mph* and *cat* operons (Fig. 3), this transcriptional cross-regulation suggests that phenol might also be degraded via *tod* enzymes, including toluene dioxygenase and the catechol *meta*-cleavage pathway (Fig. 5E). To verify the functional contributions of these genes, we constructed gene disruption mutants targeting *todE* and *mphN* in the Tol 5REK strain (Δ*todE* and Δ*mphN*), in addition to the previously constructed Δ*todC1* mutant (Yoshimoto et al., 2025). Tol 5REK carries mutations in the restriction-modification system and the *ataA* gene, enabling easier genetic manipulation and reducing cell adhesion, which is necessary for accurate growth evaluation. The *todC1* and *todE* encode the large subunit of the toluene dioxygenase complex and catechol 2,3-dioxygenase, respectively, while *mphN* encodes the large subunit of the phenol monooxygenase complex. We first confirmed none of these mutations affected growth ability in lactate culture (Fig. S7). In the toluene culture, the disruption of *todC1* or *todE* abolished growth, while the *mphN* disruption had no effect on growth (Fig. 5F). These results confirm that toluene degradation proceeds exclusively through the TOD pathway and that the phenol monooxygenase has no significant contribution to toluene assimilation. In the phenol culture, the Δ*mphN* mutant completely abolished growth, supporting the idea that phenol monooxygenase plays a major, and likely essential, role in phenol hydroxylation (Fig. 5F). In contrast, the Δ*todC1* mutant exhibited an extended lag phase, indicating that toluene dioxygenase facilitates rapid entry into exponential growth of the Tol 5 wild type on phenol. Notably, the Δ*todC1* mutant also reached a higher final cell yield compared to Tol 5REK, suggesting that toluene dioxygenase activity reduces the efficiency of carbon incorporation from phenol into biomass. The Δ*todE* mutant showed no significant difference in growth compared to Tol 5REK, suggesting that the contribution of *tod* genes to phenol assimilation operates at the hydroxylation step via toluene dioxygenase rather than through the downstream *meta*-cleavage pathway.

## 4 DISCUSSION

Members of the genus *Acinetobacter* inhabit diverse environments, including soil, oceans, freshwater, sediments, activated sludge, and polluted sites (Mateo-Estrada et al., 2019; Zhao et al., 2023). Several *Acinetobacter* strains, such as the *A. calcoaceticus–A. baumannii* complex, are also opportunistic nosocomial pathogens that exhibit multidrug resistance (Vijayakumar et al., 2019). Consequently, most previous research on *Acinetobacter* species has focused on antibiotic resistance, genomic plasticity, horizontal gene transfer, and pathogenicity, whereas the metabolic capabilities of this genus remain comparatively underexplored (Mateo-Estrada et al., 2019; Zhao et al., 2023).

Although *A. baylyi* ADP1 is a widely used model strain and its 3.5 Mb genome has been well characterized (Barbe et al., 2004; Durot et al., 2008), this genome is smaller than those of many other *Acinetobacter* species, and thus represents only a subset of the metabolic potential present across the genus (Barbe et al., 2004; Zhao et al., 2023). In this context, we characterized the carbon metabolism of the highly adhesive hydrocarbon-degrading *Acinetobacter* strain Tol 5, which has a 4.8 Mb genome (Ishikawa and Hori, 2021), providing an opportunity to explore metabolic capabilities that are underrepresented in current model strains. Genome analyses revealed that Tol 5 has limited capability to metabolize sugars but possesses an extensive repertoire of genes for alkane and aromatic metabolism. Tol 5 encodes multiple types of hydrocarbon oxygenases, including alkane monooxygenase, aromatic oxygenase, and cytochrome P450, which allow this strain to assimilate a wide range of hydrocarbons and their oxidized derivatives. Notably, we also identified five distinct aromatic degradation routes, including pathways that are rare within the genus *Acinetobacter* such as the TOD and HMG pathways, indicating metabolic versatility comparable to that of *P. putida* strains, which are widely used chassis for aromatic compound bioconversion (Nelson et al., 2002; Nogales et al., 2017; Martínez-García and de Lorenzo, 2024). These findings expand our understanding of the metabolic diversity within the genus *Acinetobacter* and highlight the utility of Tol 5 as a microbial chassis for bioproduction.

### Acinetobacter

species widely exhibit the capacity for alkane degradation, as more than 95% of strains encode the alkane monooxygenases AlkB and AlmA (Zhao et al., 2023). Oil-degrading strains such as *A. venetianus* RAG-1 and *Acinetobacter oleivorans* DR1 possess multiple AlkB homologs, enabling efficient utilization of *n*-alkanes (Park et al., 2017; Liu et al., 2021). Although Tol 5 possesses only one *alkB* gene on the genome, it also carries a plasmid-encoded cytochrome P450 monooxygenase of the CYP153 family. CYP153, which is a distinct alkane hydroxylase targeting short-and medium-chain *n*-alkanes, is typically found in alkane-degrading bacteria lacking AlkB but is relatively rare among *Acinetobacter* genomes (van Beilen Jan et al., 2006; Nie et al., 2014). Both *alkB* and the CYP153 gene were induced in Tol 5 grown on hexadecane, indicating that Tol 5 employs both types of alkane hydroxylases to oxidize hexadecane efficiently (Fig. 4D).

Regarding aromatic metabolism in *Acinetobacter* species, two main *ortho*-cleavage pathways of catechols, encoded by *pca* and *cat* genes, have been mainly studied (Barbe et al., 2004; Fischer et al., 2008; Stuani et al., 2014; Breisch et al., 2022). Typically, the *cat* operon is located adjacent to the *ben* operon, and their expression is coordinately regulated to prevent the accumulation of catechol, which is toxic at high concentration (Jiménez et al., 2014). Specifically, catechol 1,2-dioxygenase, encoded by *catA* within the ben operon, converts the catechol generated by benzoate oxidation into *cis*,*cis*-muconate, which subsequently activates the downstream *cat* operon (Cosper Nathaniel et al., 2000; Bundy et al., 2002; Silva-Rocha and de Lorenzo, 2012). In Tol 5, however, the *cat* operon is located adjacent to the *mph* operon instead of the *ben* operon (Fig. 2). Furthermore, unlike the *mph* operon of *A. calcoaceticus* PHEA-2 (Yu et al., 2011), the *mph* operon of Tol 5 contains an additional *catA* gene encoding CatAmph (*catA_2*) immediately downstream within the same operon (Fig. S3). Analogous to the coordinated regulation seen in typical *ben-cat* clusters, the presence of the *catA* gene within the *mph* operon can contribute to proper coordination with the downstream *cat* operon, ensuring the smooth conversion of the catechol derived from phenol without toxic accumulation. In addition, Tol 5 possesses a separate *ben* operon containing CatAben (*catA_4*) at a different chromosomal locus (Fig. 2). Therefore, the unique organization of the *mph* and *ben* operons in Tol 5 likely enables fine-tuned transcription of the genes involved in the degradation of phenol and probably benzoate, through the coordinated expression of CatA paralogs and their downstream catabolic operons.

Aromatic compounds with alkyl substituents are typically degraded via the *meta*-cleavage pathway, which has mainly been studied in *P. putida* (Zylstra et al., 1988; Assinder and Williams, 1990; Lacal et al., 2006). Tol 5 likely acquired these *tod* genes through horizontal gene transfer, making this strain unique within the genus *Acinetobacter* (Yoshimoto et al., 2025). In the TOD pathway, the genes encoding all reaction steps for toluene degradation, from the initial oxidation catalyzed by toluene dioxygenase to the generation of pyruvate and acetyl-CoA, are clustered in a single *tod* operon (Fig. 2). Transcriptome analysis of Tol 5 revealed that growth on toluene strongly induced the *tod* operon, whereas the *mph* operon was slightly induced (Fig. 5C). This regulatory response is consistent with the need to avoid the *ortho-*cleavage of alkyl-substituted catechols derived from toluene, which would otherwise yield dead-end metabolites such as methylmuconolactone (Taeger et al., 1988). Thus, the highly selective induction of the *tod* operon during growth on toluene suggests a transcriptional regulatory strategy that efficiently channels toluene-derived intermediates through the *meta-*cleavage pathway while minimizing flux into the unproductive *ortho-*cleavage route.

In contrast to the response to toluene, Tol 5 exhibited a distinct transcriptional pattern during growth on phenol. Phenol induced not only the *cat* and *mph* genes but also the *tod* genes (Fig. 5C). The disruption of *todC1*, encoding the large subunit of toluene dioxygenase, resulted in an extended lag phase and a higher final cell yield on phenol (Fig. 6). Previous studies have demonstrated that the toluene dioxygenase complex TodABC1C2 from *P. putida* oxidizes phenol to both catechol and hydroquinone (Höring et al., 2016). These observations suggest that, in Tol 5, the upregulation of toluene dioxygenase facilitates rapid entry into exponential growth by promoting the hydroxylation of toxic phenol, but simultaneously reduces carbon assimilation efficiency by diverting a portion of phenol into hydroquinone, for which no assimilation route was identified in Tol 5. In support of this interpretation, a mutant lacking phenol monooxygenase (Δ*mphN*) completely failed to grow on phenol (Fig. 6), indicating that the conversion of phenol by toluene dioxygenase alone is insufficient for its utilization as a carbon source. Taken together, these results indicate that phenol monooxygenase is the primary route for phenol metabolism in Tol 5, while toluene dioxygenase functions as a rapid detoxification mechanism rather than an assimilatory pathway. This stands in contrast to *P. putida* strains harboring *tod* genes, such as F1, DOT-T1, and CE2010, where the toluene dioxygenase has been recognized as the primary enzyme for phenol degradation (Spain et al., 1989; Mosqueda et al., 1999; Ohta et al., 2001; Höring et al., 2016). Our results therefore suggest a fundamentally different physiological role for the *tod* genes in Tol 5 compared with these *P. putida* strains.

Beyond the catabolic pathway for each carbon source, transcriptome analysis revealed that the number of DEGs varied substantially across conditions, with ethanol culture inducing only 14 upregulated genes compared to 266, 349, and 643 in hexadecane, toluene, and phenol cultures, respectively (Fig. 4A and 5A). This difference likely reflects the degree of metabolic remodeling required to assimilate each substrate: ethanol is converted to acetyl-CoA in only two oxidation steps and enters the TCA cycle directly, whereas hexadecane and aromatic compounds require more extensive peripheral catabolic machinery prior to entering central metabolism. Furthermore, ribosomal genes were prominently enriched among downregulated DEGs in hexadecane, toluene, and phenol cultures, suggesting a broad reduction in translational activity during growth on these substrates. Transcriptional changes were also observed in genes associated with stress response and signal transduction, including two-component systems, quorum sensing, and ABC transporters. Hydrocarbons and aromatic compounds have been reported to impose multiple types of cellular stress, such as membrane damage, the generation of toxic metabolic intermediates, and the leakage of reactive oxygen species (ROS) during their oxidation (Heipieper and Martínez, 2010; Phale et al., 2026), and the observed transcriptional changes in these categories likely represent adaptive responses to such stress conditions. However, further transcriptome analyses under more controlled conditions, such as equalized growth rates across carbon sources and reduced cell adhesion, will be required to dissect these responses in greater detail.

In conclusion, this study provides a comprehensive characterization of the carbon metabolism of *Acinetobacter* sp. Tol 5, revealing an extensive repertoire of metabolic pathways for alkanes and aromatic compounds that expands the known metabolic diversity of the genus *Acinetobacter*. Transcriptome analysis and gene disruption experiments identified the specific genes responsible for substrate assimilation and demonstrated that toluene dioxygenase and phenol monooxygenase play distinct physiological roles in phenol metabolism. These findings provide a comprehensive view of the carbon metabolism of Tol 5 and highlight its potential as a microbial chassis for bioprocesses utilizing non-sugar feedstocks.

## 5 Data availability statement

RNA-seq data reported are available in the DDBJ Sequenced Read Archive under the accession numbers PRJDB35996.

## 6 Author contributions

**Shori Inoue:** Conceptualization, Formal analysis, Investigation, Data curation, Writing - Original Draft. **Shogo Yoshimoto:** Conceptualization, Writing - Review & Editing. **Maiko Hattori:** Investigation. **Shotaro Yamagishi:** Investigation. **Katsutoshi Hori:** Conceptualization, Supervision, Project administration, Writing - Review & Editing.

## 9 Acknowledgements

The authors wish to acknowledge the Center for Gene Research, Nagoya University, for technical support with the RNA sequencing. The authors also thank Yuki Ohara for kindly discussing genome sequence data analysis and reviewing the manuscript.

## References

Anburajan, P., Naresh Kumar, A., Sabapathy, P.C., Kim, G.-B., Cayetano, R.D., Yoon, J.-J., et al. (2019). Polyhydroxy butyrate production by *Acinetobacter junii* BP25, *Aeromonas hydrophila* ATCC 7966, and their co-culture using a feast and famine strategy. Bioresource Technology 293, 122062. doi: 10.1016/j.biortech.2019.122062.

Arvay, E., Biggs, B.W., Guerrero, L., Jiang, V., and Tyo, K. (2021). Engineering *Acinetobacter baylyi* ADP1 for mevalonate production from lignin-derived aromatic compounds. Metabolic Engineering Communications 13, e00173. doi: 10.1016/j.mec.2021.e00173.

Assinder, S.J., and Williams, P.A. (1990). “The TOL Plasmids: Determinants of the Catabolism of Toluene and the Xylenes,” in Advances in Microbial Physiology, eds. A.H. Rose & D.W. Tempest. Academic Press), 1–69.

Barbe, V., Vallenet, D., Fonknechten, N., Kreimeyer, A., Oztas, S., Labarre, L., et al. (2004). Unique features revealed by the genome sequence of *Acinetobacter* sp. ADP1, a versatile and naturally transformation competent bacterium. Nucleic Acids Research 32(19), 5766–5779. doi: 10.1093/nar/gkh910.

Biggs, B.W., Bedore, S.R., Arvay, E., Huang, S., Subramanian, H., McIntyre, E.A., et al. (2020). Development of a genetic toolset for the highly engineerable and metabolically versatile *Acinetobacter baylyi* ADP1. Nucleic Acids Research 48(9), 5169–5182. doi: 10.1093/nar/gkaa167.

Bleichrodt, F.S., Fischer, R., and Gerischer, U.C. (2010). The β-ketoadipate pathway of *Acinetobacter baylyi* undergoes carbon catabolite repression, cross-regulation and vertical regulation, and is affected by Crc. Microbiology 156(5), 1313–1322. doi: 10.1099/mic.0.037424-0.

Breisch, J., Huber, L.S., Kraiczy, P., Hubloher, J., and Averhoff, B. (2022). The ß-ketoadipate pathway of *Acinetobacter baumannii* is involved in complement resistance and affects resistance against aromatic antibiotics. Environmental Microbiology Reports 14(1), 170–178. doi: 10.1111/1758-2229.13042.

Bundy, B.M., Collier, L.S., Hoover, T.R., and Neidle, E.L. (2002). Synergistic transcriptional activation by one regulatory protein in response to two metabolites. Proceedings of the National Academy of Sciences 99(11), 7693–7698. doi: 10.1073/pnas.102605799.

Chen, S., Cao, L., Lv, T., Liu, J., Gao, G., Li, M., et al. (2024). Regulation mechanism of the long-chain n-alkane monooxygenase gene *almA* in *Acinetobacter venetianus* RAG-1. Applied and Environmental Microbiology 91(1), e02050–02024. doi: 10.1128/aem.02050-24.

Chen, Y., Liu, Y., Meng, Y., Jiang, Y., Zhang, X., Liu, H., et al. (2025). Systems Metabolic Engineering of Genome-Reduced *Pseudomonas putida* for Efficient Production of Polyhydroxyalkanoate from p-Coumaric Acid. Journal of Agricultural and Food Chemistry 73(21), 12899–12907. doi: 10.1021/acs.jafc.5c02123.

Cosper Nathaniel, J., Collier Lauren, S., Clark Todd, J., Scott Robert, A., and Neidle Ellen, L. (2000). Mutations in *catB*, the Gene Encoding Muconate Cycloisomerase, Activate Transcription of the Distalben Genes and Contribute to a Complex Regulatory Circuit in *Acinetobacter* sp. Strain ADP1. Journal of Bacteriology 182(24), 7044–7052. doi: 10.1128/jb.182.24.7044-7052.2000.

D’Almeida, A.P., de Azevedo, D.C.S., Melo, V.M.M., de Albuquerque, T.L., and Rocha, M.V.P. (2024). Bioemulsifier production by *Acinetobacter venetianus* AMO1502: Potential for bioremediation and environmentally friendly applications. Marine Pollution Bulletin 203, 116436. doi: 10.1016/j.marpolbul.2024.116436.

Dal, S., Trautwein, G., and Gerischer, U. (2005). Transcriptional Organization of Genes for Protocatechuate and Quinate Degradation from *Acinetobacter* sp. Strain ADP1. Applied and Environmental Microbiology 71(2), 1025–1034. doi: 10.1128/AEM.71.2.1025-1034.2005.

de Lorenzo, V., Pérez-Pantoja, D., and Nikel Pablo, I. (2024). *Pseudomonas putida* KT2440: the long journey of a soil-dweller to become a synthetic biology chassis. Journal of Bacteriology 206(7), e00136–00124. doi: 10.1128/jb.00136-24.

Durot, M., Le Fèvre, F., de Berardinis, V., Kreimeyer, A., Vallenet, D., Combe, C., et al. (2008). Iterative reconstruction of a global metabolic model of *Acinetobacter baylyi* ADP1 using high-throughput growth phenotype and gene essentiality data. BMC Systems Biology 2(1), 85. doi: 10.1186/1752-0509-2-85.

Fischer, R., Bleichrodt, F.S., and Gerischer, U.C. (2008). Aromatic degradative pathways in *Acinetobacter baylyi* underlie carbon catabolite repression. Microbiology 154(10), 3095–3103. doi: 10.1099/mic.0.2008/016907-0.

Heipieper, H.J., and Martínez, P.M. (2010). “Toxicity of Hydrocarbons to Microorganisms,” in Handbook of Hydrocarbon and Lipid Microbiology, ed. K.N. Timmis. (Berlin, Heidelberg: Springer Berlin Heidelberg), 1563–1573.

Hooppaw Anna, J., McGuffey Jenna, C., Di Venanzio, G., Ortiz-Marquez Juan, C., Weber Brent, S., Lightly Tasia, J., et al. (2022). The Phenylacetic Acid Catabolic Pathway Regulates Antibiotic and Oxidative Stress Responses in *Acinetobacter*. mBio 13(3), e01863–01821. doi: 10.1128/mbio.01863-21.

Hori, K., Ishikawa, M., Yamada, M., Higuchi, A., Ishikawa, Y., and Ebi, H. (2011). Production of peritrichate bacterionanofibers and their proteinaceous components by *Acinetobacter* sp. Tol 5 cells affected by growth substrates. Journal of Bioscience and Bioengineering 111(1), 31–36. doi: 10.1016/j.jbiosc.2010.08.009.

Hori, K., Yamashita, S., Ishii, S.i., Kitagawa, M., Tanji, Y., and Unno, H. (2001). Isolation, Characterization and Application to Off-Gas Treatment of Toluene-Degrading Bacteria. Journal of Chemical Engineering of Japan 34(9), 1120–1126.

Höring, P., Rothschild-Mancinelli, K., Sharma, N.D., Boyd, D.R., and Allen, C.C.R. (2016). Oxidative biotransformations of phenol substrates catalysed by toluene dioxygenase: A molecular docking study. Journal of Molecular Catalysis B: Enzymatic 134, 396–406. doi: 10.1016/j.molcatb.2016.10.013.

Inoue, S., Yoshimoto, S., and Hori, K. (2025). A new target of multiple lysine methylation in bacteria. Journal of Bacteriology 207(1), e00325–00324. doi: 10.1128/jb.00325-24.

Ishige, T., Tani, A., Sakai, Y., and Kato, N. (2000). Long-Chain Aldehyde Dehydrogenase That Participates in n-Alkane Utilization and Wax Ester Synthesis in *Acinetobacter* sp. Strain M-1. Applied and Environmental Microbiology 66(8), 3481–3486. doi: 10.1128/AEM.66.8.3481-3486.2000.

Ishikawa, M., and Hori, K. (2021). Complete Genome Sequence of the Highly Adhesive Bacterium *Acinetobacter* sp. Strain Tol 5. Microbiology Resource Announcements 10(35), doi: 10.1128/mra.00567-21.

Ishikawa, M., and Hori, K. (2024). The elimination of two restriction enzyme genes allows for electroporation-based transformation and CRISPR-Cas9-based base editing in the non-competent Gram-negative bacterium *Acinetobacter* sp. Tol 5. Applied and Environmental Microbiology 90(6), e00400-00424. doi: 10.1128/aem.00400-24.

Ishikawa, M., Nakatani, H., and Hori, K. (2012a). AtaA, a New Member of the Trimeric Autotransporter Adhesins from *Acinetobacter* sp. Tol 5 Mediating High Adhesiveness to Various Abiotic Surfaces. PLOS ONE 7(11), e48830. doi: 10.1371/journal.pone.0048830.

Ishikawa, M., Shigemori, K., Suzuki, A., and Hori, K. (2012b). Evaluation of adhesiveness of *Acinetobacter* sp. Tol 5 to abiotic surfaces. Journal of Bioscience and Bioengineering 113(6), 719–725. doi: 10.1016/j.jbiosc.2012.01.011.

Jiménez, J.I., Pérez-Pantoja, D., Chavarría, M., Díaz, E., and de Lorenzo, V. (2014). A second chromosomal copy of the *catA* gene endows *Pseudomonas putida* mt-2 with an enzymatic safety valve for excess of catechol. Environmental Microbiology 16(6), 1767–1778. doi: 10.1111/1462-2920.12361.

Kong, W., Zhao, C., Gao, X., Wang, L., Tian, Q., Liu, Y., et al. (2021). Characterization and Transcriptome Analysis of a Long-Chain n-Alkane-Degrading Strain *Acinetobacter pittii* SW-1. International Journal of Environmental Research and Public Health 18(12), 6365.

Kurnia, K., Efimova, E., Santala, V., and Santala, S. (2024). Metabolic engineering of *Acinetobacter baylyi* ADP1 for naringenin production. Metabolic Engineering Communications 19, e00249. doi: 10.1016/j.mec.2024.e00249.

Lacal, J., Busch, A., Guazzaroni, M.-E., Krell, T., and Ramos, J.L. (2006). The TodS–TodT two-component regulatory system recognizes a wide range of effectors and works with DNA-bending proteins. Proceedings of the National Academy of Sciences 103(21), 8191–8196. doi: 10.1073/pnas.0602902103.

Lin, G.-H., Hsieh, M.-C., and Shu, H.-Y. (2021). Role of Iron-Containing Alcohol Dehydrogenases in *Acinetobacter baumannii* ATCC 19606 Stress Resistance and Virulence. International Journal of Molecular Sciences 22(18), 9921.

Liu, H., Tao, X., Ntakirutimana, S., Liu, Z.-H., Li, B.-Z., and Yuan, Y.-J. (2024). Engineering *Pseudomonas putida* for lignin bioconversion into *cis*-*cis* muconic acid. Chemical Engineering Journal 495, 153375. doi: 10.1016/j.cej.2024.153375.

Liu, J., Zhao, B., Lan, Y., and Ma, T. (2021). Enhanced degradation of different crude oils by defined engineered consortia of *Acinetobacter venetianus* RAG-1 mutants based on their alkane metabolism. Bioresource Technology 327, 124787. doi: 10.1016/j.biortech.2021.124787.

Luo, J., McIntyre Emily, A., Bedore Stacy, R., Santala, V., Neidle Ellen, L., and Santala, S. (2022). Characterization of Highly Ferulate-Tolerant *Acinetobacter baylyi* ADP1 Isolates by a Rapid Reverse Engineering Method. Applied and Environmental Microbiology 88(2), e01780–01721. doi: 10.1128/AEM.01780-21.

Maier, T., Förster, H.-H., Asperger, O., and Hahn, U. (2001). Molecular Characterization of the 56-kDa CYP153 from *Acinetobacter* sp. EB104. Biochemical and Biophysical Research Communications 286(3), 652–658. doi: 10.1006/bbrc.2001.5449.

Martínez-García, E., and de Lorenzo, V. (2024). *Pseudomonas putida* as a synthetic biology chassis and a metabolic engineering platform. Current Opinion in Biotechnology 85, 103025. doi: 10.1016/j.copbio.2023.103025.

Mateo-Estrada, V., Graña-Miraglia, L., López-Leal, G., and Castillo-Ramírez, S. (2019). Phylogenomics Reveals Clear Cases of Misclassification and Genus-Wide Phylogenetic Markers for *Acinetobacter*. Genome Biology and Evolution 11(9), 2531–2541. doi: 10.1093/gbe/evz178.

McClure, R., Balasubramanian, D., Sun, Y., Bobrovskyy, M., Sumby, P., Genco, C.A., et al. (2013). Computational analysis of bacterial RNA-Seq data. Nucleic Acids Research 41(14), e140–e140. doi: 10.1093/nar/gkt444.

Mosqueda, G., Ramos-González, M.a.-I., and Ramos, J.L. (1999). Toluene metabolism by the solvent-tolerant *Pseudomonas putida* DOT-T1 strain, and its role in solvent impermeabilization. Gene 232(1), 69–76. doi: 10.1016/S0378-1119(99)00113-4.

Nakazawa, T. (2002). Travels of a *Pseudomonas*, from Japan around the world. Environmental Microbiology 4(12), 782–786. doi: 10.1046/j.1462-2920.2002.00310.x.

Nelson, K.E., Weinel, C., Paulsen, I.T., Dodson, R.J., Hilbert, H., Martins dos Santos, V.A.P., et al. (2002). Complete genome sequence and comparative analysis of the metabolically versatile *Pseudomonas putida* KT2440. Environmental Microbiology 4(12), 799–808. doi: 10.1046/j.1462-2920.2002.00366.x.

Nie, Y., Chi, C.-Q., Fang, H., Liang, J.-L., Lu, S.-L., Lai, G.-L., et al. (2014). Diverse alkane hydroxylase genes in microorganisms and environments. Scientific Reports 4(1), 4968. doi: 10.1038/srep04968.

Nogales, J., García, J.L., and Díaz, E. (2017). “Degradation of Aromatic Compounds in *Pseudomonas*: A Systems Biology View,” in Aerobic Utilization of Hydrocarbons, Oils and Lipids, ed. F. Rojo. Springer, Cham), 1–49.

Ohara, Y., Yoshimoto, S., and Hori, K. (2019). Control of AtaA-mediated bacterial immobilization by casein hydrolysates. Journal of Bioscience and Bioengineering 128(5), 544–550. doi: 10.1016/j.jbiosc.2019.04.019.

Ohta, Y., Maeda, M., and Kudo, T. (2001). *Pseudomonas putida* CE2010 can degrade biphenyl by a mosaic pathway encoded by the *tod* operon and *cmtE*, which are identical to those of *P. putida* F1 except for a single base difference in the operator–promoter region of the *cmt* operon. Microbiology 147(1), 31–41. doi: 10.1099/00221287-147-1-31.

Park, C., Shin, B., Jung, J., Lee, Y., and Park, W. (2017). Metabolic and stress responses of *Acinetobacter oleivorans* DR1 during long-chain alkane degradation. Microbial Biotechnology 10(6), 1809–1823. doi: 10.1111/1751-7915.12852.

Phale, P.S., Dhamale, T., and Subhash, S. (2026). The Arsenal of Aromatic Degrading Bacteria: How They Sense, Chase, Adapt and Destroy Environmental Pollutants. Microbial Biotechnology 19(5), e70372. doi: 10.1111/1751-7915.70372.

Phale, P.S., Malhotra, H., and Shah, B.A. (2020). “Chapter One - Degradation strategies and associated regulatory mechanisms/features for aromatic compound metabolism in bacteria,” in Advances in Applied Microbiology, eds. G.M. Gadd & S. Sariaslani. Academic Press), 1–65.

Pontrelli, S., Chiu, T.-Y., Lan, E.I., Chen, F.Y.H., Chang, P., and Liao, J.C. (2018). *Escherichia coli* as a host for metabolic engineering. Metabolic Engineering 50, 16–46. doi: 10.1016/j.ymben.2018.04.008.

Ramos, J.-L., Sol Cuenca, M., Molina-Santiago, C., Segura, A., Duque, E., Gómez-García, M.R., et al. (2015). Mechanisms of solvent resistance mediated by interplay of cellular factors in *Pseudomonas putida*. FEMS Microbiology Reviews 39(4), 555–566. doi: 10.1093/femsre/fuv006.

Ren, X., and Palmer Lauren, D. (2023). *Acinetobacter* Metabolism in Infection and Antimicrobial Resistance. Infection and Immunity 91(6), e00433–00422. doi: 10.1128/iai.00433-22.

Rojo, F. (2009). Degradation of alkanes by bacteria. Environmental Microbiology 11(10), 2477–2490. doi: 10.1111/j.1462-2920.2009.01948.x.

Ruhl, I.A., Woodworth, S.P., Haugen, S.J., Alt, H.M., Beckham, G.T., and Johnson, C.W. (2025). Production of Vanillin From Ferulic Acid by *Pseudomonas putida* KT2440 Using Metabolic Engineering and In Situ Product Recovery. Microbial Biotechnology 18(5), e70152. doi: 10.1111/1751-7915.70152.

Seo, P.-W., Hwangbo, S.-A., Kim, J.-S., and Park, S.-Y. (2025). Structural Mimicry Without Glyoxalase I Functional Convergence: A Homogentisate 1,2-Dioxygenase From *Acinetobacter*. Proteins: Structure, Function, and Bioinformatics 93(12), 2150–2157. doi: 10.1002/prot.70020.

Silva-Rocha, R., and de Lorenzo, V. (2012). A GFP-*lacZ* Bicistronic Reporter System for Promoter Analysis in Environmental Gram-Negative Bacteria. PLOS ONE 7(4), e34675. doi: 10.1371/journal.pone.0034675.

Spain, J.C., Zylstra, G.J., Blake, C.K., and Gibson, D.T. (1989). Monohydroxylation of phenol and 2,5-dichlorophenol by toluene dioxygenase in *Pseudomonas putida* F1. Applied and Environmental Microbiology 55(10), 2648–2652. doi: 10.1128/aem.55.10.2648-2652.1989.

Stuani, L., Lechaplais, C., Salminen, A.V., Ségurens, B., Durot, M., Castelli, V., et al. (2014). Novel metabolic features in *Acinetobacter baylyi* ADP1 revealed by a multiomics approach. Metabolomics 10(6), 1223–1238. doi: 10.1007/s11306-014-0662-x.

Taboada, B., Estrada, K., Ciria, R., and Merino, E. (2018). Operon-mapper: a web server for precise operon identification in bacterial and archaeal genomes. Bioinformatics 34(23), 4118–4120. doi: 10.1093/bioinformatics/bty496.

Taeger, K., Knackmuss, H.-J., and Schmidt, E. (1988). Biodegradability of mixtures of chloro-and methylsubstituted aromatics: Simultaneous degradation of 3-chlorobenzoate and 3-methylbenzoate. Applied Microbiology and Biotechnology 28(6), 603–608. doi: 10.1007/BF00250420.

Teufel, R., Mascaraque, V., Ismail, W., Voss, M., Perera, J., Eisenreich, W., et al. (2010). Bacterial phenylalanine and phenylacetate catabolic pathway revealed. Proceedings of the National Academy of Sciences 107(32), 14390–14395. doi: 10.1073/pnas.1005399107.

Usami, A., Ishikawa, M., and Hori, K. (2018). Heterologous expression of geraniol dehydrogenase for identifying the metabolic pathways involved in the biotransformation of citral by *Acinetobacter* sp. Tol 5. Bioscience, Biotechnology, and Biochemistry 82(11), 2012–2020. doi: 10.1080/09168451.2018.1501263.

Usami, A., Ishikawa, M., and Hori, K. (2020). Gas-phase bioproduction of a high-value-added monoterpenoid (*E*)-geranic acid by metabolically engineered *Acinetobacter* sp. Tol 5. Green Chemistry 22(4), 1258–1268. doi: 10.1039/C9GC03478A.

van Beilen Jan, B., Funhoff Enrico, G., van Loon, A., Just, A., Kaysser, L., Bouza, M., et al. (2006). Cytochrome P450 Alkane Hydroxylases of the CYP153 Family Are Common in Alkane-Degrading Eubacteria Lacking Integral Membrane Alkane Hydroxylases. Applied and Environmental Microbiology 72(1), 59–65. doi: 10.1128/AEM.72.1.59-65.2006.

Vijayakumar, S., Indranil, B., and Veeraraghavan, B. (2019). Accurate Identification of Clinically Important *Acinetobacter* Spp.: An Update. Future Science OA 5(6), FSO395. doi: 10.2144/fsoa-2018-0127.

Wang, Y., Wang, Z., Chen, Y., Hua, X., Yu, Y., and Ji, Q. (2019). A Highly Efficient CRISPR-Cas9-Based Genome Engineering Platform in *Acinetobacter baumannii* to Understand the H2O2-Sensing Mechanism of OxyR. Cell Chemical Biology 26(12), 1732–1742.e1735. doi: 10.1016/j.chembiol.2019.09.003.

Wohlfarth, S., Hoesche, C., Strunk, C., and Winkler, U.K. (1992). Molecular genetics of the extracellular lipase of *Pseudomonas aeruginosa* PAO1. Microbiology 138(7), 1325–1335. doi: 10.1099/00221287-138-7-1325.

Yoshimoto, S., Aoki, S., Ohara, Y., Ishikawa, M., Suzuki, A., Linke, D., et al. (2023). Identification of the adhesive domain of AtaA from *Acinetobacter* sp. Tol 5 and its application in immobilizing *Escherichia coli*. Frontiers in Bioengineering and Biotechnology 10(1095057). doi: 10.3389/fbioe.2022.1095057.

Yoshimoto, S., Hattori, M., Inoue, S., Mori, S., Ohara, Y., and Hori, K. (2025). Identification of toluene degradation genes in *Acinetobacter* sp. Tol 5. Journal of Bioscience and Bioengineering 140, 284-289. doi: 10.1016/j.jbiosc.2025.07.010.

Yoshimoto, S., Ishii, S., Kawashiri, A., Matsushita, T., Linke, D., Göttig, S., et al. (2024). Adhesion preference of the sticky bacterium *Acinetobacter* sp. Tol 5. Frontiers in Bioengineering and Biotechnology 12, 1342418. doi: 10.3389/fbioe.2024.1342418.

Yoshimoto, S., Ohara, Y., Nakatani, H., and Hori, K. (2017). Reversible bacterial immobilization based on the salt-dependent adhesion of the bacterionanofiber protein AtaA. Microbial Cell Factories 16(1), 123. doi: 10.1186/s12934-017-0740-7.

Yu, G., Wang, L.-G., Han, Y., and He, Q.-Y. (2012). clusterProfiler: an R Package for Comparing Biological Themes Among Gene Clusters. OMICS: A Journal of Integrative Biology 16(5), 284–287. doi: 10.1089/omi.2011.0118.

Yu, H., Peng, Z., Zhan, Y., Wang, J., Yan, Y., Chen, M., et al. (2011). Novel Regulator MphX Represses Activation of Phenol Hydroxylase Genes Caused by a XylR/DmpR-Type Regulator MphR in *Acinetobacter calcoaceticus*. PLOS ONE 6(3), e17350. doi: 10.1371/journal.pone.0017350.

Zhao, Y., Wei, H.-M., Yuan, J.-L., Xu, L., and Sun, J.-Q. (2023). A comprehensive genomic analysis provides insights on the high environmental adaptability of *Acinetobacter* strains. Frontiers in Microbiology 14, 1177951. doi: 10.3389/fmicb.2023.1177951.

Zylstra, G.J., McCombie, W.R., Gibson, D.T., and Finette, B.A. (1988). Toluene degradation by *Pseudomonas putida* F1: genetic organization of the tod operon. Applied and Environmental Microbiology 54(6), 1498–1503. doi: 10.1128/aem.54.6.1498-1503.1988.

