## Supplementary Material for "Characterization of carbon metabolism in a highly adhesive bacterium *Acinetobacter* sp. Tol 5 capable of assimilating diverse hydrocarbons and aromatic compounds"

### Supplementary Figures

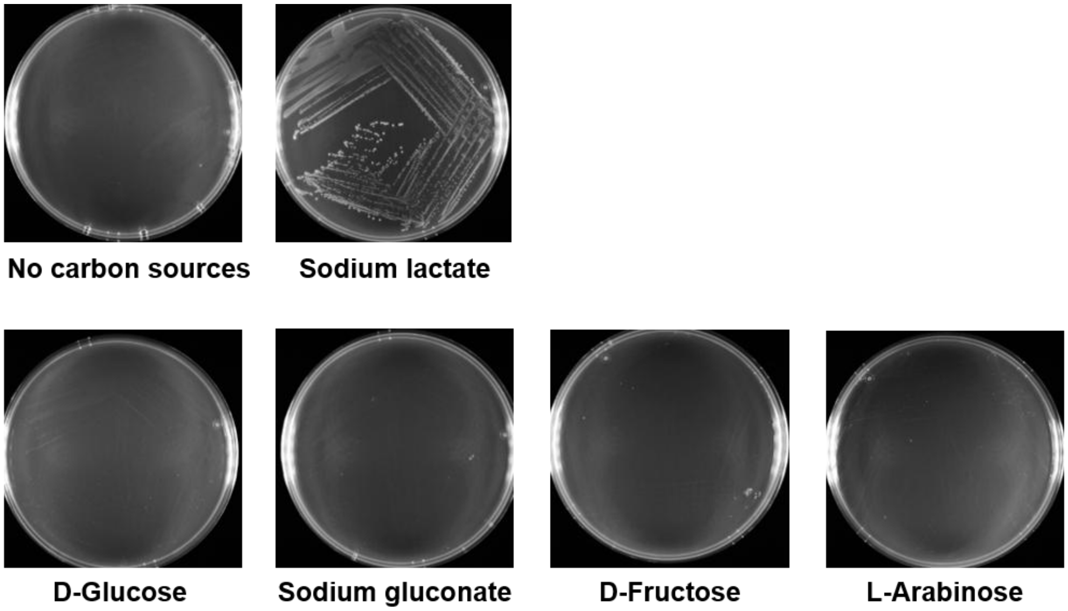


**Supplementary Figure 1.** Growth of Tol 5 on sugars. The Tol 5 Δ*ataA* mutant was streaked on BS agar plates supplemented with sodium lactate, D-glucose, sodium gluconate, D-fructose, or L-arabinose as the sole carbon source and incubated for 5 days.

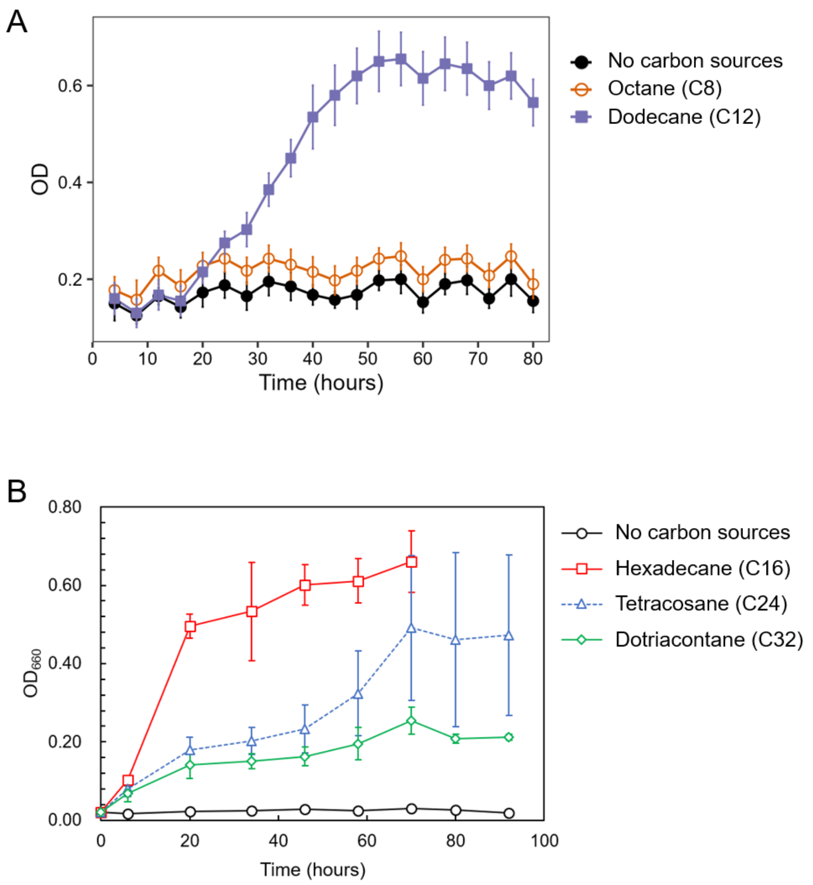


**Supplementary Figure 2.** Growth of Tol 5 on *n*-alkanes of different chain lengths. The Tol 5 Δ*ataA* mutant was grown in BS medium with each *n*-alkane as the sole carbon source. (**A**) Growth on octane (C8) and dodecane (C12), measured in glass tubes with automatic OD monitoring using the OD-monitor C&T system. (**B**) Growth on hexadecane (C16), tetracosane (C24), and dotriacontane (C32), measured in flasks by sampling and measuring $OD_{660}$ with the UV–Vis spectrophotometer. All data are presented as the means ± SEMs (biological replicates n = 3).

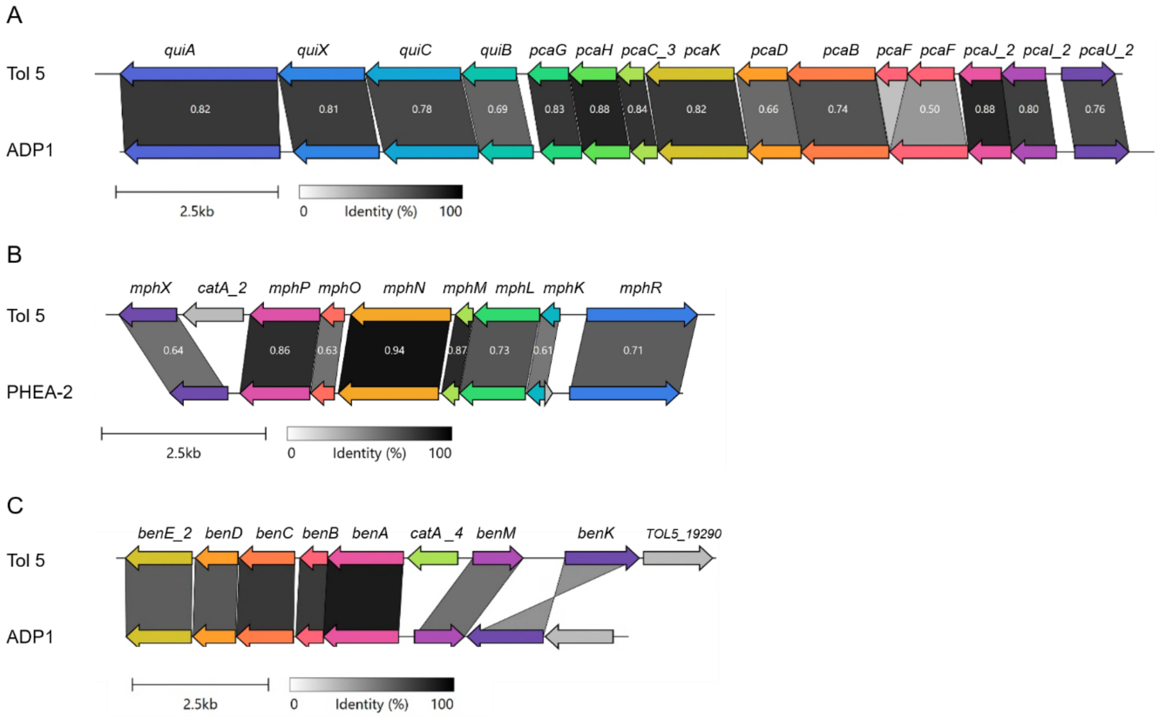


**Supplementary Figure 3.** Organization of aromatic degradation gene clusters in Tol 5 compared with related *Acinetobacter* species. Schematics represent the genomic organization of the *pca-qui* (**A**), *mph* (**B**), and *ben* (**C**) gene clusters of Tol 5, aligned using clinker with the corresponding clusters of *A. baylyi ADP1* (accession: CR543861.1) or *A. calcoaceticus* PHEA-2 (accession: CP002177.1). Arrows indicate predicted coding sequences and their transcriptional orientation. Homologous genes are connected between clusters.

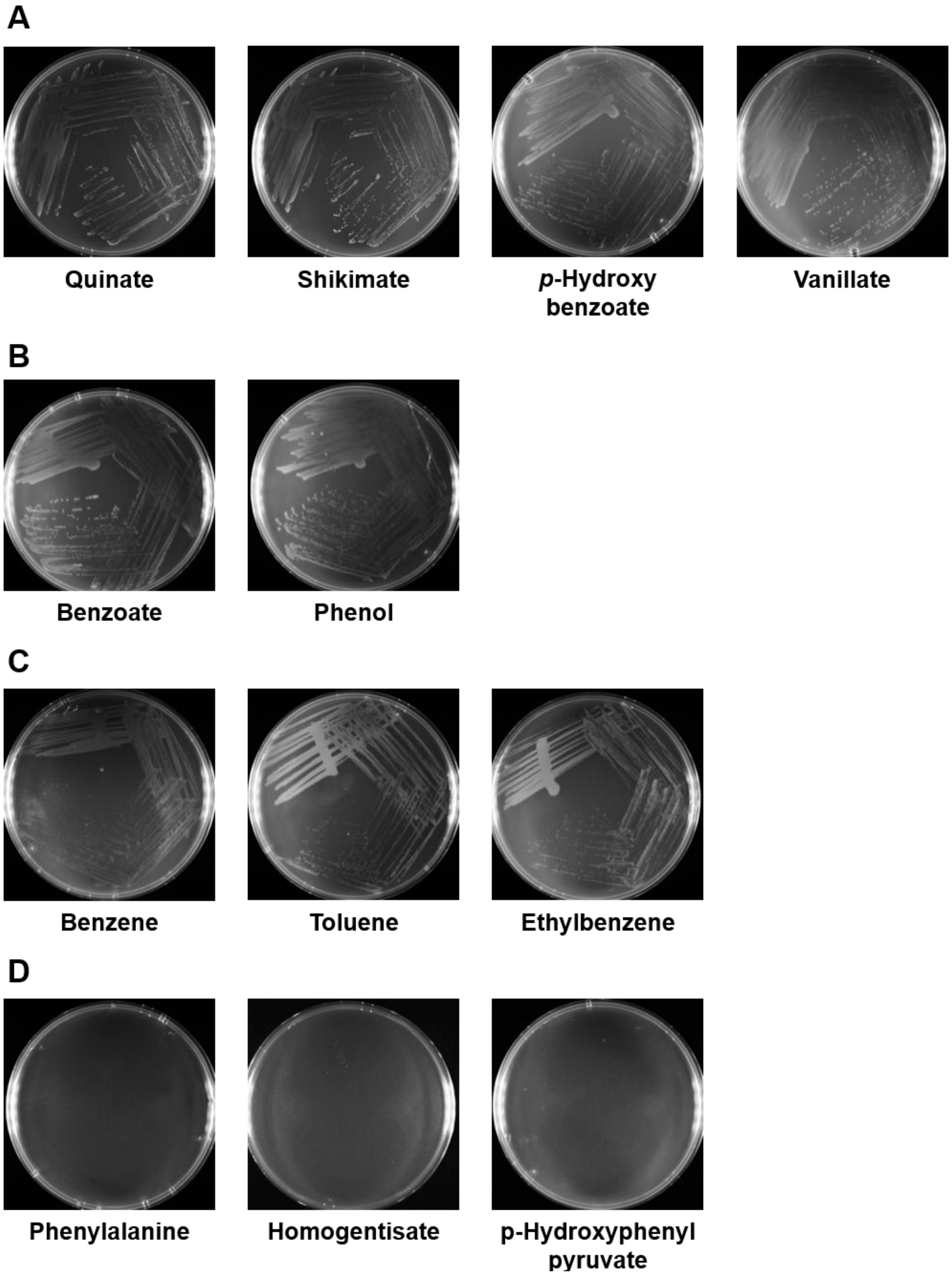


**Supplementary Figure 4.** Growth of Tol 5 on aromatic compounds on agar plates. The Tol 5 ΔataA mutant was streaked on BS agar plates supplemented with aromatic compound as the sole carbon source and incubated for 5 days. Plates were sealed in AnaeroPack pouches and incubated for 5 days. Volatile substrates were supplied in the vapor phase. (**A**) Substrates related to the PCA pathway: quinate, shikimate, *p*-hydroxybenzoate, and vanillate. (**B**) Substrates related to the CAT pathway: phenol and benzoate. (**C**) Substrates related to the TOD pathway: toluene, benzene, and ethylbenzene. (**D**) Substrates related to aromatic amino acid metabolism: L-phenylalanine, homogentisate, and *p*-hydroxyphenylpyruvate.

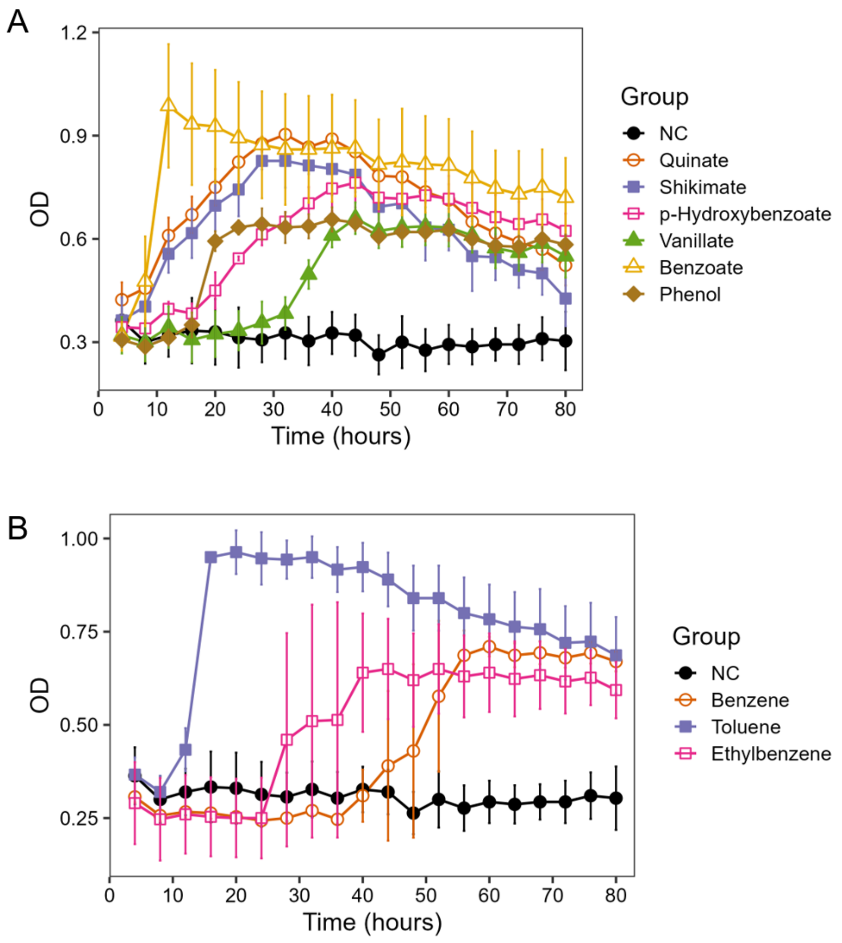


**Supplementary Figure 5.** Growth of Tol 5 on aromatic substrates. The Tol 5 Δ*ataA* mutant was grown in BS medium with each aromatic substrate as the sole carbon source, and the OD was automatically monitored in glass tubes using the OD-monitor C&T system. (**A**) Substrates funneled through the PCA and CAT pathways: quinate, shikimate, *p*-hydroxybenzoate, vanillate, benzoate, and phenol. (**B**) Substrates degraded via the TOD pathway: benzene, toluene, and ethylbenzene. NC, no carbon source control. Data are presented as the means ± SEM (n = 3 biological replicates).

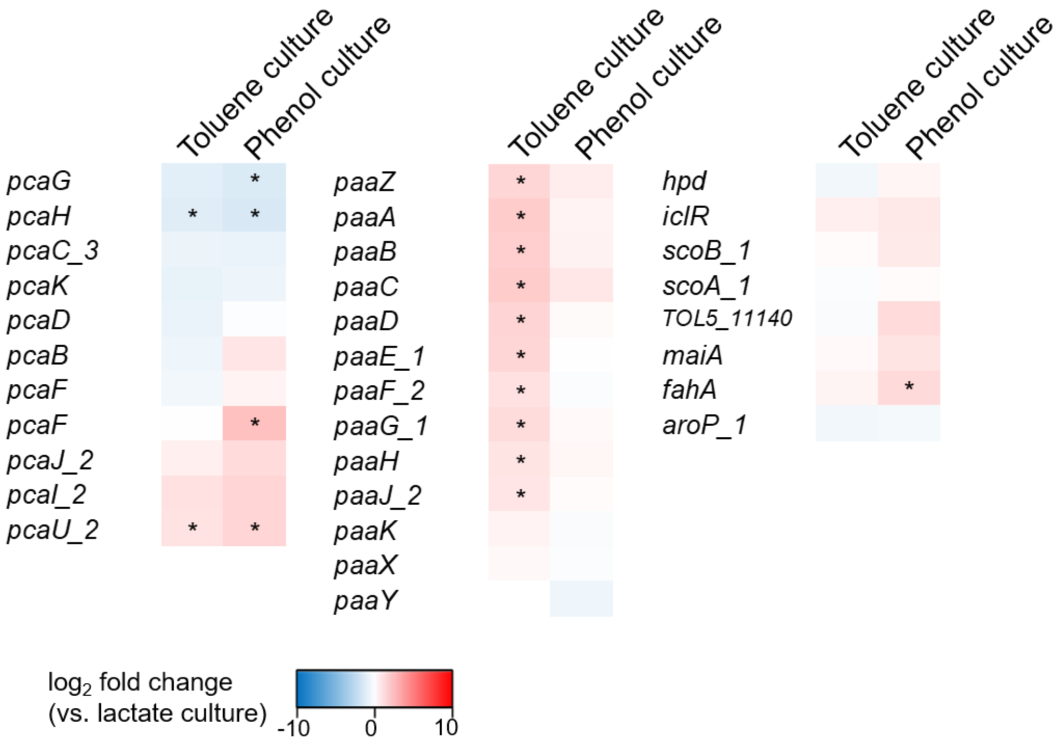


**Supplementary Figure 6.** Heat maps showing the average $\log_2$ FC of the genes involved in the degradation of aromatic compounds (*pca*, *paa*, and *hmg* genes). The color gradient indicates the magnitude of the $\log_2$ FC values, with the deepest red representing ≥ 10 and the deepest blue representing ≤ -10. Asterisks (*) denote differentially expressed genes defined by $|\log_2 \text{FC}| > 1$, $\log_2$ CPM > 3, and FDR < 0.01.

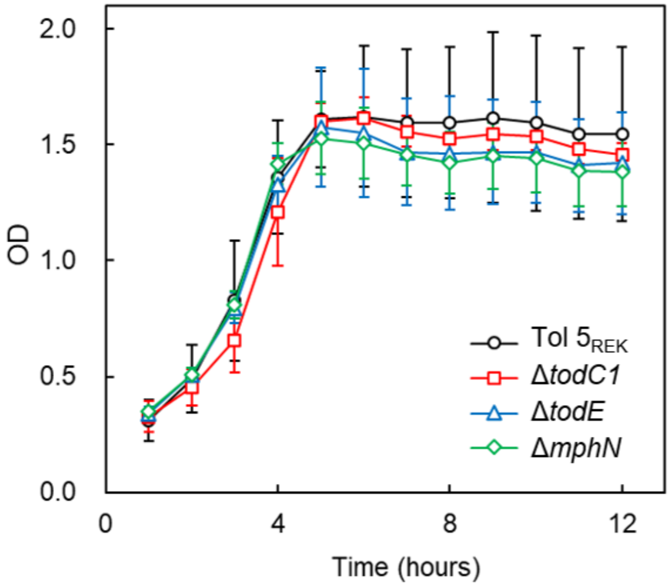


**Supplementary Figure 7.** Growth of gene disruption mutants on sodium lactate. Tol $5_{REK}$ (circles), the Δ*todC1* mutant (squares), the Δ*todE* mutant (triangles), and the Δ*mphN* mutant (diamonds) were inoculated to BS medium containing lactate at a carbon equivalent concentration of $1.4 \times 10^{-2}$ mol/L, and OD was monitored using the OD-monitorC&T system. Data are presented as the means ± SEMs (biological replicates n = 3).

**Supplemental Tables**

**Supplementary Table 1.** Strains and plasmids used in this study.

| Strain | Description | Reference |
|---|---|---|
| *Acinetobacter* sp. | | |
| Tol 5 | Wild type strain | (Hori et al., 2001) |
| Tol 5 Δ*ataA* | *ataA* deletion mutant of Tol 5 | (Ishikawa et al., 2012) |
| Tol $5_{REK}$ | Restriction enzyme-encoding genes and *ataA* knockout mutant of Tol 5, REK123Δ*ataA* | (Ishikawa and Hori, 2024) |
| Tol 5 $_{REK}$ Δ*todC1* | *todC1* gene knockout mutant of Tol $5_{REK}$ | (Yoshimoto et al., 2025) |
| Tol 5 $_{REK}$ Δ*todE* | *todE* gene knockout mutant of Tol $5_{REK}$ | This study |
| Tol $5_{REK}$ Δ*mphN* | *mphN* gene knockout mutant of Tol $5_{REK}$ | This study |
| *Escherichia coli* | | |
| DH5α | Host for routine cloning | Purchased from Takara Bio (Shiga, Japan) |
| Plasmid | | |
| pBECAb-apr | For gene knockout | (Wang et al., 2019) |
| pBECAb-apr-todE | For *todE* gene knockout | This study |
| pBECAb-apr-mphN | For *mphN* gene knockout | This study |

**Supplementary Table 2.** Oligo DNAs used in this study.

| Primer name | Sequence (5' to 3') |
|---|---|
| TodE-sgRNA-Fw | CTAAGCGGTCTCTTAGTAATCGAGTTGAGGTGTAT TAGTTTGGAGACCCGTAGC |
| TodE-sgRNA-Rv | GCTACGGGTCTCCAAACTAATACACCTCAACTCGA TTACTAAGAGACCGCTTAG |
| MphN-sgRNA-Fw | TAGTAAACTCCCAGTCCAGATCAC |
| MphN-sgRNA-Rv | AAACGTGATCTGGACTGGGAGTTT |

**Supplementary Table 3.** Functional predictions of genes in Tol 5 based on multiple databases. (Excel file)

**Supplementary Table 4**. Genes predicted to be involved in the metabolism in Tol 5. (Excel file)

**Supplementary Table 5.** Information about the RNA-seq data used in this study

| Substrates | Total reads (raw FASTQ) | Total reads after fastp filtering | Total mapped reads | Mapping rate (%) |
|---|---|---|---|---|
| Lactate | 12813627 | 12653268 | 11336668 | 89.6 |
| | 12421215 | 12298185 | 10803072 | 87.8 |
| | 12339378 | 12221016 | 11190080 | 91.6 |
| Ethanol | 12814610 | 12700050 | 11516082 | 90.7 |
| | 11870987 | 11766340 | 10751266 | 91.4 |
| | 12887700 | 12771057 | 11582603 | 90.7 |
| Hexadecane | 11487878 | 11231169 | 9913352 | 88.3 |
| | 11814155 | 11698621 | 10507217 | 89.8 |
| | 12704470 | 12510943 | 11381040 | 91.0 |
| Toluene | 12476544 | 12235527 | 11115768 | 90.8 |
| | 11862557 | 11754526 | 10750795 | 91.5 |
| | 10735004 | 10650101 | 9637121 | 90.5 |
| Phenol | 11282222 | 11144094 | 10095402 | 90.6 |
| | 10571014 | 10447506 | 9312043 | 89.1 |
| | 10544517 | 10445993 | 9367032 | 89.7 |

**Supplementary Table 6.** Comparison of gene expression in Tol 5 grown on various carbon sources relative to lactate. (Excel file)

**Supplementary Table 7.** Differentially expressed genes in Tol 5 grown on various carbon sources relative to lactate. (Excel file)

**References**


Hori, K., Yamashita, S., Ishii, S.i., Kitagawa, M., Tanji, Y., and Unno, H. (2001). Isolation, Characterization and Application to Off-Gas Treatment of Toluene-Degrading Bacteria. *Journal of Chemical Engineering of Japan* 34(9)**,** 1120-1126.

Ishikawa, M., and Hori, K. (2024). The elimination of two restriction enzyme genes allows for electroporation-based transformation and CRISPR-Cas9-based base editing in the non-competent Gram-negative bacterium *Acinetobacter* sp. Tol 5. *Applied and Environmental Microbiology* 90(6)**,** e00400-00424. doi: 10.1128/aem.00400-24.

Ishikawa, M., Nakatani, H., and Hori, K. (2012). AtaA, a New Member of the Trimeric Autotransporter Adhesins from *Acinetobacter* sp. Tol 5 Mediating High Adhesiveness to Various Abiotic Surfaces. *PLOS ONE* 7(11)**,** e48830. doi: 10.1371/journal.pone.0048830.

Wang, Y., Wang, Z., Chen, Y., Hua, X., Yu, Y., and Ji, Q. (2019). A Highly Efficient CRISPR-Cas9-Based Genome Engineering Platform in *Acinetobacter baumannii* to Understand the $H_2O_2$-Sensing Mechanism of OxyR. *Cell Chemical Biology* 26(12)**,** 1732-1742.e1735. doi: https://doi.org/10.1016/j.chembiol.2019.09.003.

Yoshimoto, S., Hattori, M., Inoue, S., Mori, S., Ohara, Y., and Hori, K. (2025). Identification of toluene degradation genes in *Acinetobacter* sp. Tol 5. *Journal of Bioscience and Bioengineering* 140**,** 284-289. doi: 10.1101/2025.05.12.653411.
